# Were Neoarchean atmospheric methane hazes and early Paleoproterozoic glaciations driven by the rise of oxygen in surface environments?

**DOI:** 10.1101/2020.10.30.362202

**Authors:** Anshuman Swain, Alan J Kaufman, Marcin Kalinowski, Stephanie A Yarwood, William F Fagan

## Abstract

Geochemical evidence suggests methane was the predominant greenhouse gas in the Archean and early Proterozoic eons. Consequently, fluctuations in methane concentration in Earth’s atmosphere (i.e., ‘methane hazes’) would have contributed to climate change and influenced the flux of UV radiation reaching surface environments. If correct, understanding what factors (e.g., O_2_ and/or resource concentration) drove the biological methane cycle might shed light on the repetition of biological, atmospheric and climatic events preserved in the sedimentary rock record from ~2.8 to 2.0 BYA. To explore these interdependencies, we developed a novel dynamical model of microbial ecological interactions to investigate the conditions under which methane is preferentially released to the atmosphere. We found that the interplay between resource and O_2_ availability results in complex cyclic methane dynamics unrelated to the functional groups or efficiencies of microbial communities, to initial conditions, or to other model constraints. Based on these results, we propose that the cyclicity of methane haze events and glacial episodes in the Neoarchean and early Paleoproterozoic may have been linked to the progressive increase in oceanic and atmospheric O_2_ through the interval.

## Introduction

Like Titan, Saturn’s orange-shrouded moon, the early Earth may have been episodically enveloped in a thick blanket of atmospheric methane (CH_4_), a powerful greenhouse gas produced by archaeal methanogens in anoxic environments. A methane haze would have formed if the CH_4_/CO_2_ ratio was >0.2 (Trainer et al., 2004; Domagal-Goldman et al., 2008) contrasting with the value of 0.005 in the modern oxygenated atmosphere. Although the depth and distribution of the CH_4_ haze during such events likely varied across the planet due to vertical atmospheric motion and mixing (Wolf and Toon, 2010), the biogenic gas would nonetheless have warmed the troposphere, and through attenuation of the solar UV flux, protected surface environments from high energy radiation.

Carbon isotope evidence from Neoarchean (2.8 to 2.5 BYA) organic matter preserved in sedimentary rocks (Eigenbrode and Freeman, 2006) suggests a prolonged increase in the production of CH_4_ and its consumption by methanotrophs to form biomass. Depending on the leak rate of the greenhouse gas from anoxic sediments, CH_4_ may have accumulated in sufficient concentrations in the atmosphere to form periodic hazes. The extent and depth of such hazes is likely to have been modulated by the photosynthetic production and accumulation of molecular oxygen (O_2_) in the oceans and atmosphere.

Due to the rapid oxidation of CH_4_ to CO_2_ in the presence of O_2_ (Saunois et al., 2016), the atmospheric concentration of CH_4_ is generally thought to represent a balance between source and sink reactions. Tipping the scales between the two would thus modulate the size of the atmospheric reservoir of CH_4_. On the source side, the production of CH_4_ could have been affected by competition for critical metabolic substrates between methanogens and sulfate reducers, or by the lack of key metal cofactors, like nickel, required for methanogenic enzymes (Anbar and Knoll, 2002; Konhauser et al., 2009). In this analysis, however, we consider ecological factors and biological interactions that may have increased the CH_4_ flux from surface environments that are directly associated with the release of photosynthetic O_2_. Such a scenario would be particularly relevant in the Neoarchean when periodic CH_4_ hazes are suggested from geochemical measurements of sedimentary rocks and photochemical models (Zerkle et al., 2012; Farquhar et al., 2013; Izon et al., 2015; 2017, Williford et at., 2016).

Our dynamic resource-driven model for CH_4_ generation and release is based upon the metabolic repercussions of the Warburg Effect in microbial functional groups (see Swain and Fagan, 2018) – which was initially proposed to explain uncontrolled cellular proliferation during cancer growth (Warburg, 1956) – to an ecological scenario appropriate to Neoarchean and early Paleoproterozoic surface environments and atmospheric fluctuations. The Warburg Effect in microorganisms describes a shift in metabolic respiration where the energetic process of oxidative phosphorylation (OP), which produces CO_2_ as an end product, is replaced by glycolysis that instead generates products that can be incorporated into biomass but yields 10 times less energy. This shift in respiration mechanisms is particularly pronounced during the rapid growth of microbial cells that are not resource limited – even in the presence of oxygen. This metabolic phenomenon has been observed in a wide variety of organisms such as bacteria (where it is termed “overflow metabolism”; Basan et al., 2015, Schuster et al., 2015; Zhuang et al., 2011), yeast (the “Crabtree effect”; Deken, 1966), and a diversity of human cells, including Kupffer and microglial cells (specialized macrophages in the liver and brain, respectively), lymphocytes (white blood cells), and fast twitch muscle fibers (Schuster et al., 2015; Vander Heiden et al., 2009). Especially in microorganisms, this phenomenon is known to have cascading effects on the stoichiometry and community dynamics in natural ecosystems (see Manzoni et al., 2012; Carlson et al., 2018; Kreft et al., 2020). While the Warburg Effect may seem energetically counterintuitive, one can observe that as cells grow rapidly, the kinetic advantage of glycolysis that occurs in the cytoplasm results in a two-fold energy advantage over OP, which is restricted to the cell membrane (Voet and Voet, 2004). Past models thus describe a metabolic arms race between OP and glycolysis that includes: i) efficiency vs. rate (OP is more efficient, but glycolysis is faster), ii) surface area vs. volume (glycolysis occurs in the cell interior whereas OP is constrained to the cell surface), and iii) biomass production (glycolysis produces organic substrates that can be used in biomass production directly, which is generally not true for OP) (see Swain and Fagan, 2018; Schuster et al., 2015). Results from these models suggest that large inputs of resources (i.e., any carbon substrate that allows the microbes to grow) with even small inputs of oxygen can stimulate glycolysis over the more energetic process of OP, and therefore, can affect ecological processes, specifically methanogenesis, that use these glycolytic overflow products (e.g., formate, lactate, and acetate).

To understand the dynamics of how resource and O_2_ availability can affect the production and consumption of CH_4_, we present a dynamical model of microbial ecological interactions, using the concept of overflow metabolism associated with the Warburg Effect. The ecological interactions described in this ordinary differential equation (ODE) model may be relevant to geological episodes where pulsed resource inputs or the availability of O_2_ can alter the CH_4_ production rate. Our ODE model might then apply to climatic and environmental perturbations of the early Earth, including the cyclic patterns of Neoarchean CH_4_ haze events (Pavlov et al., 2001; Zerkle et al., 2012; Izon et al., 2015; 2017) and early Paleoproterozoic glacial episodes (e.g., Bekker and Kaufman, 2007).

In our model, we focus on five major functional groups of microbes: obligate aerobic heterotrophic bacteria (A), methanogens (B), aerobic methanotrophs (C), generalized fermenters (F), anaerobic methanotrophs (G; also known as the anaerobic oxidation of methane (AOM) consortia), and anaerobic heterotrophs (H; such as iron and sulfur reducers). All of these microbial functional groups are involved in the production and/or consumption of generalized resources (R), glycolytic/fermentation products (FP), O_2_, CH_4_, and CO_2_ (**Fig. 1**). Here, the generalized resources are an abstract representation of a variety of materials that organisms might use as a carbon source, or in an indirect assimilation of carbon source and growth. Through this generalization of resources, we allow that some carbon-based compounds come from oxygenic photosynthesis. For example, anoxygenic photosynthesis (which, in turn, depended on and was limited by electron acceptors; see Ward et al., 2019) appears to have provided carbon sources for microbes, especially in Neoarchean and early Paleoproterozoic sedimentary environments. As a consequence of this generalization of resources, our model considers a decoupled system of R and O_2_. In addition, non-carbon resources can vary in their stoichiometry in organisms (see Sterner and Elser, 2002); therefore, this decoupling of R and O_2_ also helps to decode the effects of the continental nutrient influx to the oceans, or related phenomena.

**Figure 1:**
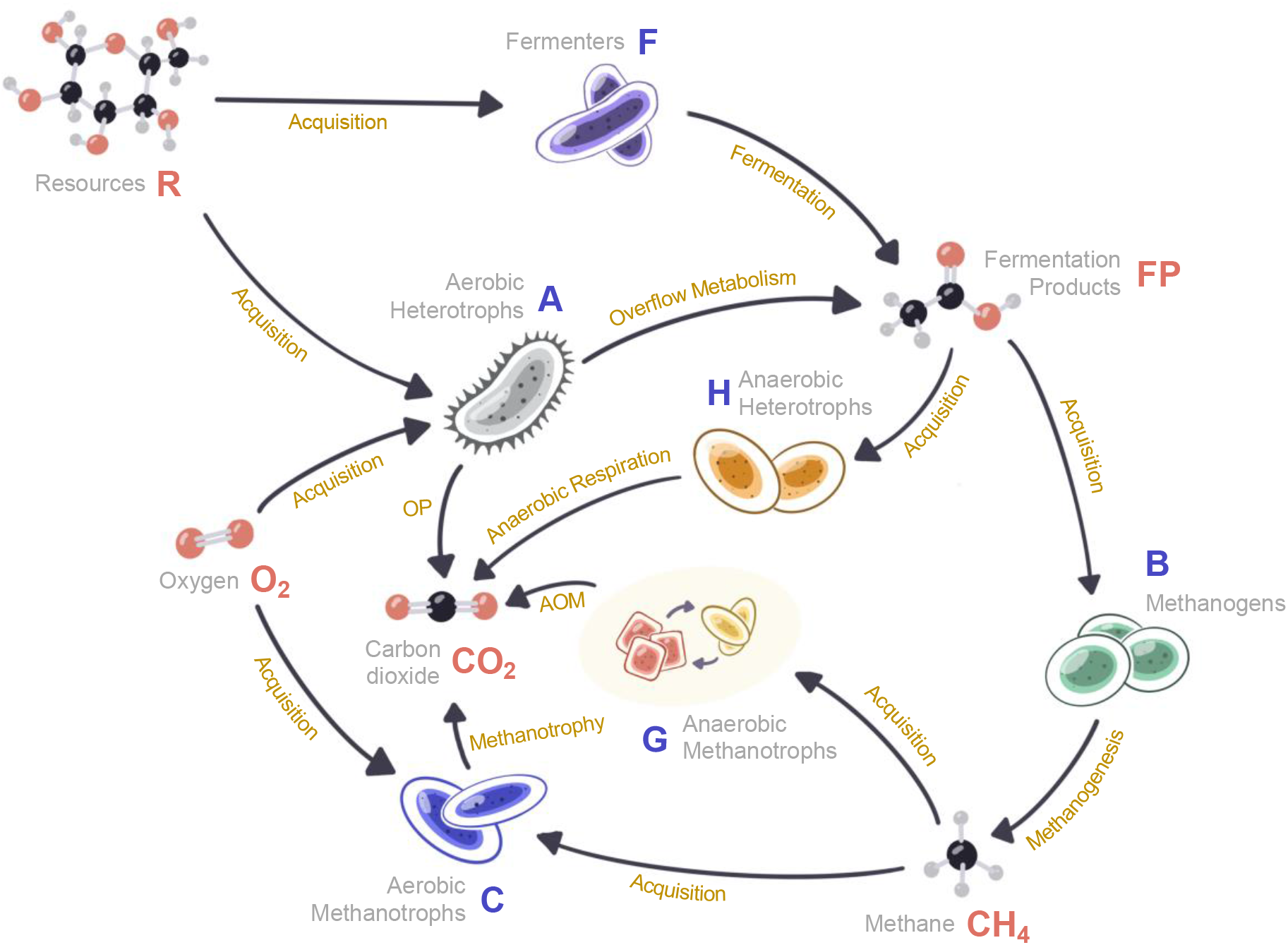
Flow diagram representation of the model system. Here (in blue) A, B, C, F, G and H are functional groups of heterotrophic bacteria, methanogens, aerobic methanotrophs, fermenters, anaerobic methanotrophs, and anaerobic heterotrophs, respectively. The flux arrows indicate processes that produce (in red) either metabolic reactants or gases cycled among the microbial functional groups. Ordinary differential equations with detailed explanations representing this dynamical system appear in **Methods**.

Key relationships within the flow diagram (Fig. 1) can be understood as follows. Methanogens depend upon fermentation products (FP) for their methane production; FP are simple organic substrates such as formate, lactate, and acetate that are produced by Fermenters (F). Aerobic heterotrophic bacteria (A) also produce fermentation products (FP) through overflow metabolism, insofar as this functional group responds to increasing R by producing more FP than CO_2_, and thereby stimulating methanogenesis (see **Methods**). Anaerobic heterotrophs (H) compete with methanogens (B) for FP. Methane is consumed by aerobic methanotrophs (C) that utilize O_2_ to produce CO_2_, and by anaerobic methanotrophs (G), in consortia with various other groups of microbes, to produce either alkalinity or CO_2_ through the anaerobic oxidation of methane (AOM). Changes in the size of microbial functional groups are modelled as functions of the availability of R and the efficiency of their utilization (energy production), as well as their basal energy requirements necessary to maintain homeostasis (energy requirement). We assume methanogens (B) represent both acetoclastic and hydrogenotrophic methanogenesis with respect to reduced FP compounds. Acetoclastic methanogens use primary FP, and hydrogenotrophic methanogens depend on secondary fermentation for H_2_ and formate. We did not include the hydrogenotrophic methanogens as a sink for CO_2_, however, because this can cause unintended model estimation errors that are important for the decoupling between R and O_2_. As we are assuming a decoupling between CO_2_, O_2_, and R (given the unknown extent of oxygenic photosynthesis), we can either leave CO_2_ as a sink (as modeled) or couple CO_2_ back to R as a function of oxygenic and anoxygenic autotrophy. In the latter case, we would include an explicit CO_2_ flux to the hydrogenotrophic methanogens. In the former (as chosen), we assume only that there is enough CO_2_ to fulfill all necessary demands, and therefore the substrate is not a limiting reagent.

The main rationale behind constructing this coupled ODE-based ecological model is to understand the response of microbial functional groups to the availability of R and O_2_, through mutual interactions and usage of substrates. Changes in the abundance of resources, metabolic products (FP, CH_4_, and CO_2_), and O_2_ associated with consortia interactions are functions of efficiencies of i) overflow metabolism, ii) respiration, and iii) biomass production and maintenance, as well as the rate of addition of both resources and O_2_. To explore the model, we considered both driven and non-driven states. First, for non-driven dynamics, we explore the basic dependence of metabolic parameters and species interactions in closed systems wherein microbes and resources start from an initial condition, and are then closed off to external changes. Second, we consider driven dynamics in open systems, in which there are constant rates of addition of R and O_2_. Numerical exploration of the model suggest that the interplay between the two results in complex, but robust, cyclic methane dynamics. We relate this new dynamic perspective to the episodic nature of Neoarchean methane haze events and early Paleoproterozoic glaciation.

## Results

### No external forcing

For the case of no external forcing (non-driven) where the only initial inputs were R and O_2_, we ran 500,000 model simulations to steady-state (**Figure S1**). Across these simulations, we varied key model parameters (the efficiency of various metabolic reactions, including methanogenesis, AOM, and fermentation) and initial conditions (starting populations of different microbial functional groups, and original abundance of R and O_2_) using Latin Hypercube Sampling (LHS; Carnell, 2020) to explore their impacts on final gas (O_2_, CO_2_, and CH_4_) concentrations. We gauged the importance of the different parameters and conditions using partial rank correlation coefficients (PRCC) (see **Methods**), and found that steady-state CH_4_ and CO_2_ concentrations varied the most based upon the metabolic parameters of methanogens (these included resource incorporation efficiency (*β*), resource utilization efficiency (*κ*) and energy requirement (*u*)) (**Figure S2B,C**). The model further reveals that steady state CO_2_ concentration varied strongly as a function of the metabolic parameters that controlled anaerobic methanotrophy (or AOM, G) (**Figure S2B**). Steady state O_2_ concentrations varied as functions of the metabolic parameters of aerobic methanotrophs (C), aerobic heterotrophic bacteria (A), and methanogens (B) (**Figure S2A**). None of the initial conditions that we varied (i.e., starting populations of different microbial functional groups and initial concentrations of R and O_2_) had a high PRCC for any of the stable state gas concentrations **(Figure S3)**, except for the dependence of the final CO_2_ concentration on the initial value of O_2_ concentration **(Figure S3B)**.

Although the steady state gas concentrations varied quantitatively with certain parameters, the qualitative trends that they depicted were relatively invariant across the Latin Hypercube sweep of parameter space and initial conditions. **Figure 2** depicts a specific case, where we plot the steady state variations of the proportions of various gases in a closed, non-driven system (initialized with a biologically-relevant base parametrization) due to changes in the methane production efficiency coefficient (*β*, see Eqs. 4, 9, and 11 in **Methods**), which was set at 1%, 10%, and 50%. With increasing efficiency, the system became more anoxic (lower O_2_) and less acidic (lower CO_2_), while CH_4_ concentrations increased, especially at low initial O_2_ and high initial R inputs. We recorded similar model responses for variations of all other parameters (See **Figures S4A-O**). However, we noted lower than expected CO_2_ concentrations at low initial O_2_ inputs for low efficiencies of parameters governing the incorporation and utilization of resources by the aerobic heterotrophs (see Eqs. 2, 3, 8, and 10 in **Methods**; and **Figs. S4D and S4H**); these can be attributed to the PRCC dependence of CO_2_ on initial O_2_ concentration. We also noted a minor deviation in CH_4_ concentration at a high initial O_2_ input in the *Ω* (resource utilization efficiency for anaerobic methanotrophs, G; see **Methods**) simulations (see Eqs. 7 and 10 in **Methods**; **Fig. S4K**); this was attributable to the formation of CO_2_ and its dependence on metabolic parameters of G. These non-linearities arose in cases where the efficiency of usage, or of incorporation, is either biologically unrealistic, or are at the extreme end of what is possible for biological systems (e.g., a very low O_2_ affinity for aerobic heterotrophs or a high CH_4_ usage efficiency in anaerobic methanotrophs [see Morris and Schmidt 2013; Bowles et al., 2019]). Overall, these results signify that, on average, the qualitative outputs of our ODE computations remained consistent across the parametrizations and initial conditions in the non-driven setting.

**Figure 2:**
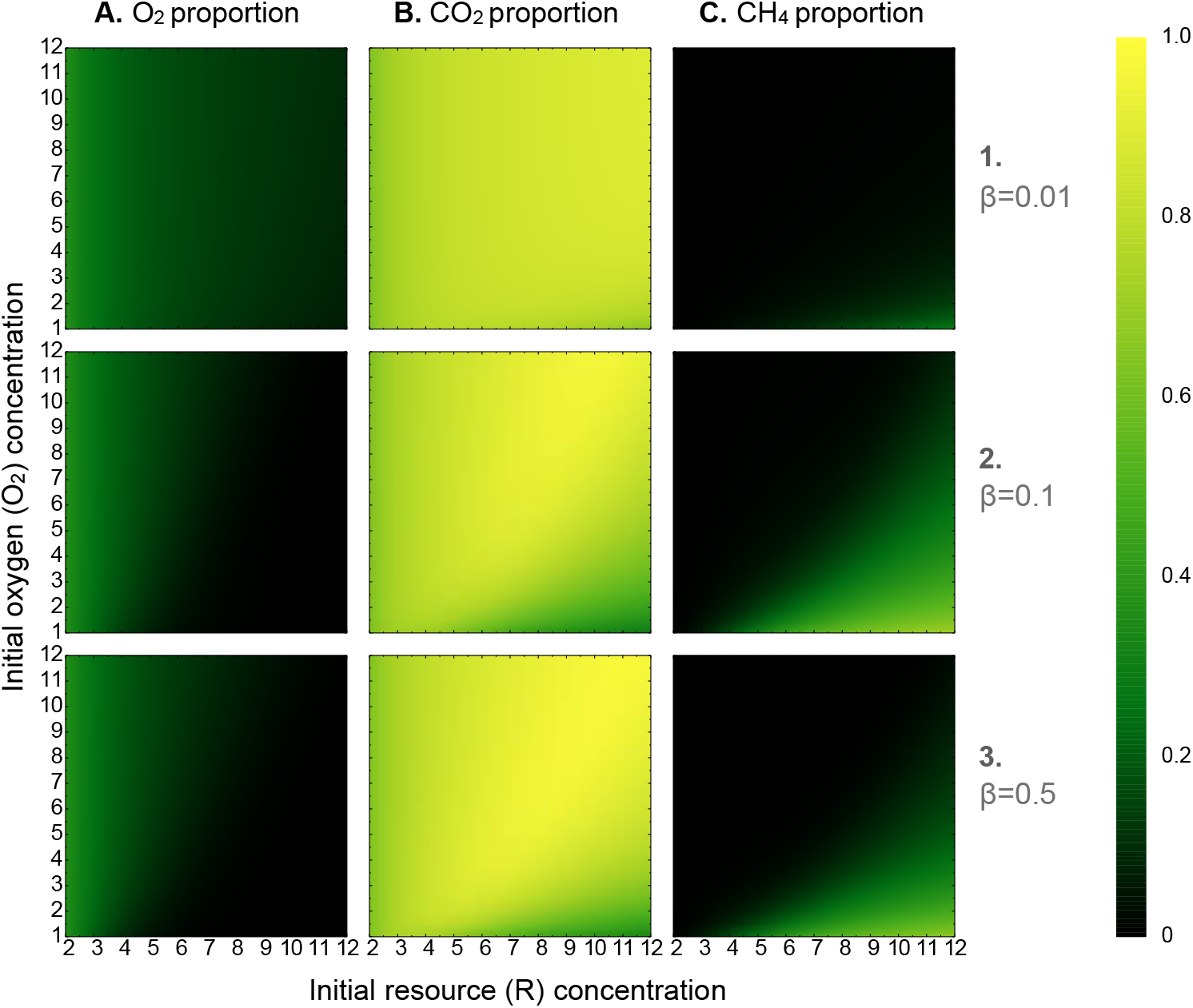
Dynamic variations in O_2_, CO_2_, and CH_4_ proportions (in columns A-C) run to steady state at set O_2_ and R input values, with the coefficient of CH_4_ production (*β*) set at 1, 10, and 50% in rows 1-3, respectively. Note that the resource (R) axis begins at 2; this is due to constraints posed by the minimal energy requirements for each microbial functional group, below which all functional groups diminish.

We further tested the stability and the sensitivity of the modeled system to the addition of new microbial functional groups to the main structure by comparing with results of a model lacking H (anaerobic heterotrophs) that compete directly with B (methanogens). The comparison shows no significant differences among the two models in terms of their partial rank correlation coefficients (see Figure S2).

### External forcing

In contrast with the no external forcing scenario above, microbial systems may also be forced, particularly by influxes of metabolizable organic substrates (resource, R) and O_2_ that are linked through oxygenic photosynthesis, wherein R is synthesized and O_2_ is released as a byproduct of the metabolic activity. Model runs with only forced R inputs were qualitatively simple. As illustrated in **Fig. S5**, initiating the model with a constant rate of addition of R depletes O_2_ rapidly regardless of its initial abundance. Under these conditions, the Warburg Effect in oxidative heterotrophs does not play a major role in the long-term dynamics of the microbial system. In this scenario, the Warburg Effect only affects the system initially, so that most FP is derived from general fermenters (F) for the subsequent production of CH_4_ through methanogenesis. Most notably, simulations with lower initial O_2_ (**Fig. S5B**) resulted in an oscillatory behavior over longer periods of CH_4_ generation. These results contrast with simulations with higher initial O_2_ (**Fig. S5A**), which resulted in rapid damping of the oscillatory behavior. Note, however, that all simulations with the same parameter values reached the same steady state irrespective of the initial conditions, as one can extrapolate from **Figure S5** (and PRCC values from steady state dynamics). Thus, these simulations revealed that differences in the initial O_2_ input only affected the relative timescale of events, as the saturation values (amplitudes) and profiles of the CH_4_, CO_2_ and R concentrations remained the same. Higher initial O_2_ stabilized the system more rapidly to steady state condition, while higher initial R under low O_2_ resulted in sustained oscillations over longer intervals. If the initial R input is held constant, the dynamic solution to forcing the microbial system with a fixed rate of addition of O_2_ is that CH_4_ will be produced briefly associated with the initial concentration of resource, and O_2_ will fall as long as methane is produced and aerobic methanotrophs consume it (**Fig. S6**). After the CH_4_ resource is depleted, the continued input of O_2_ overwhelms the system and kills the anaerobes, thereby allowing O_2_ to increase at a linear rate. Different starting concentrations of R have no apparent effect on system-level dynamics.

Truly complex dynamics occur when both R and O_2_ are supplied at constant rates (the input rate per unit time, denoted as *σ*_1_(*t*) for R and *σ*_2_(*t*) for O_2_; see Eqs. 8 and 12 in **Methods**) (**Fig. 3).** Low oxygen supply rates relative to resource supply rates (i.e., high *σ*_2_(*t*) / *σ*_1_(*t*)) or conversely low *σ*_2_(*t*) / *σ*_1_(*t*)) result in large amplitude, slow frequency oscillations in methane and oxygen concentrations in the system, a phenomenon that differs from the model behavior with only resource addition (see **Figure S5; Figure 3A**). With decreasing *σ*_2_(*t*) / *σ*_1_(*t*)), these oscillations reduce in amplitude and increase in frequency until a threshold value (of relative O_2_ input), where the system reaches a non-oscillatory stable state (**Figure 3A-J; Figure 4**). The transitions into and out of the periodic behavior appear smooth, and the amplitude and periodicity of the CH_4_ peaks within the dynamic interval decline with increasing O_2_ input rate. Moreover, the ratio of resource to oxygen input rates at which oscillatory behavior transitions to a stable, non-oscillatory pattern remains constant across the parameter variations and initial conditions (i.e. the ratio of R to O_2_ input remained at 5, or, *σ*_2_(*t*) / *σ*_1_(*t*)~5)(**Figure 4B**). **Figure 4A** further illustrates the relationship between the O_2_ input rate (*σ*_2_, varied from 10 to 20% of *σ*_1_) on the amplitude and periodicity of CH_4_ peaks. Clearly the greatest amplitudes and periods in CH_4_ concentration occur at low O_2_ input rates, and these both decline rapidly as oxygen overwhelms the system, leading to a phase transition when *σ*_2_ is ~ 20% of *σ*_1_ (**Figures 3 and 4**).

**Figure 3:**
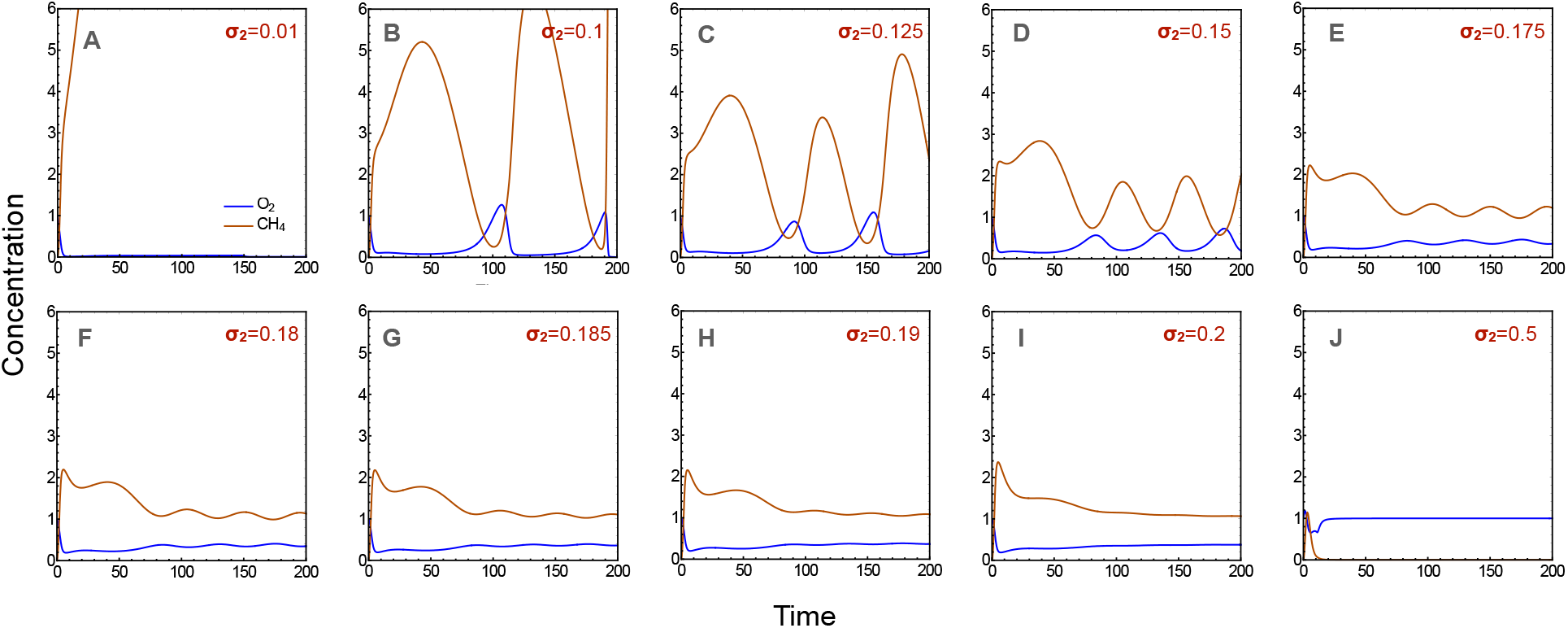
Dynamic variations in O_2_ and CH_4_ for the externally forced model with constant rates of addition of R and O_2_. The O_2_ input rate (*σ*_2_) varied between 1% to 50% relative to the resource input rate across panels (*σ*_1_), but in all cases the R input rate (*σ*_1_) was held constant at 1.

**Figure 4:**
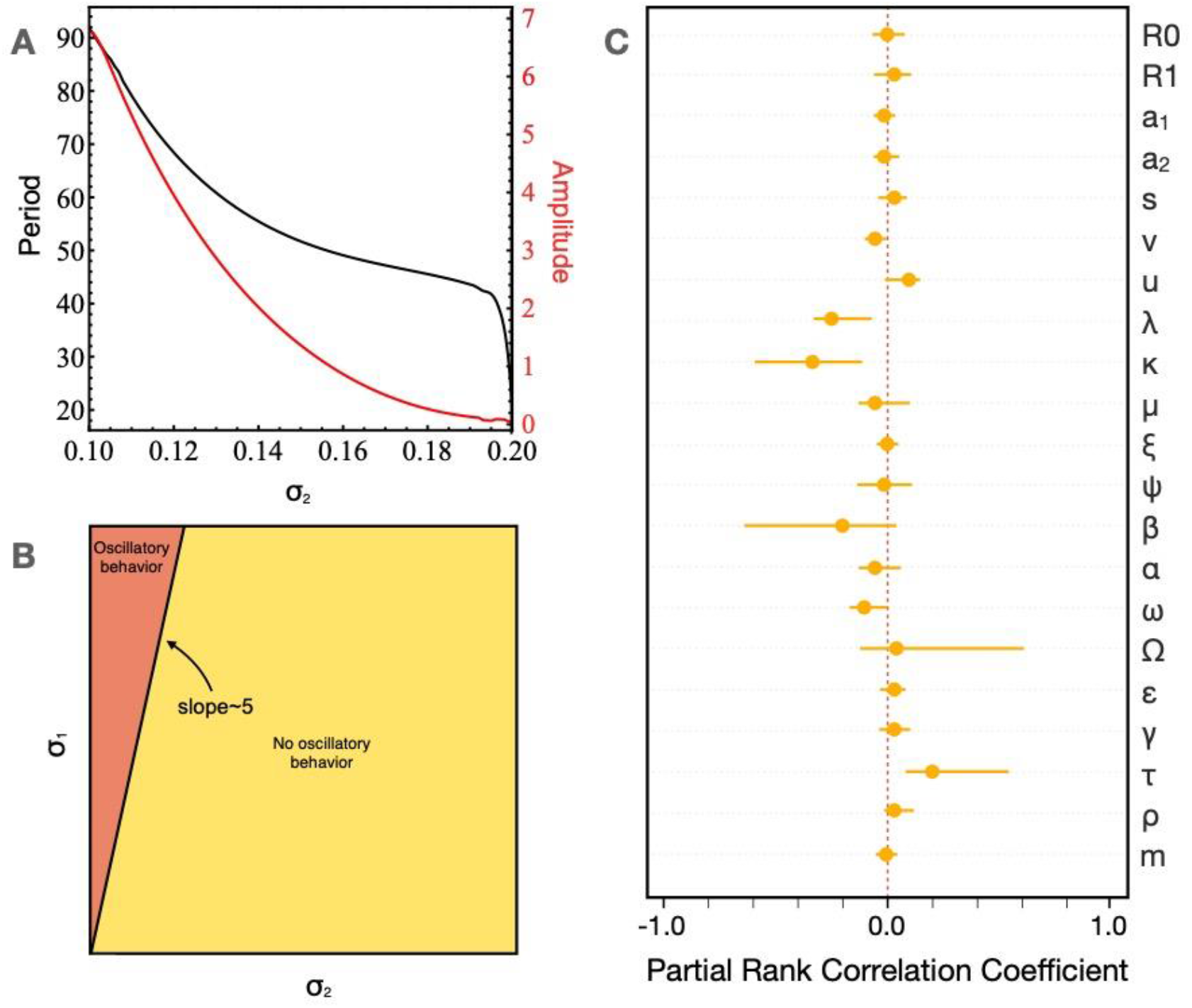
(A) shows the dependence of amplitude (in red) and time period (in black) of the CH_4_ oscillations in Fig. 3 as a function of the O_2_ input rate (*σ*_2_). (B) demarcates the resource input (*σ*_1_) -oxygen input (*σ*_2_) space on the basis of oscillatory behavior for our standard set of parameters. (C) depicts the dependence of the slope of the line separating the space between oscillatory and non-oscillatory behavior in *σ*_1_-*σ*_2_ space on specific parameters (for description of parameters, see **Methods**; Table 1) over a wide range of parameter combinations through partial rank correlation coefficient (PRCC).

We conduct a similar robustness test for the dynamics of a reduced model (lacking the H functional group of anaerobic heterotrophs) with the dynamics of the full model, and find negligible differences between the two (e.g., Figure S7). This signifies that the model is mostly affected by the relative addition rates of oxygen to resources, and not by other parameters or initial conditions (see Figure S2, S3, S7) for large qualitative changes.

We also perform a sensitivity analysis of the slope demarcating the oscillatory and non-oscillatory behavior in a variety of parameter combination (selected through LHS) with the results illustrated in Figure 4C. We find that the space demarcation is affected most by the resource utilization efficiency of methanogens (*β*), anaerobic methanotrophs (*Ω*) and anaerobic heterotrophs ( ), as well as resource incorporation efficiency of aerobic methanotrophs (*λ*) and methanogens (*κ*). Increased conversion efficiency of carbon substrates to methane (represented by *β*) has been hypothesized to result in severe biotic crises and methanogenic bursts in the past (Rothman et al., 2014). The role of aerobic and anaerobic methanotrophs (represented by *Ω* and *λ, respectively*) is usually to control the large-scale methane flux, and that of anaerobic heterotrophs (through) is to compete for carbon substrates with methanogens (Lyu et al., 2018).

## Discussion

Our dynamic, resource-driven ecological model based on the Warburg Effect reveals clues to the production of CH_4_ in response to the availability of molecular O_2_ when applied to a consortium of aerobic and anaerobic microbial functional groups. By progressively increasing the input rate of O_2_ from 10-20% with respect to the resource input rate, the overflow metabolism of the heterotrophic bacteria provides the resources necessary to produce CH_4_ in greater proportions than it is consumed by both aerobic and anaerobic methanotrophs.

Applied to potentially coupled biological and environmental events across the Neoarchean-early Paleoproterozoic transition, this dynamic ecological model may help to explain how the evolution of oxygenic photosynthesis drove fluctuations in the oxidation state of surface environments, as well as the pulsed releases of CH_4_ to the atmosphere that formed periodic methane hazes. Carbon isotope compositions of organic matter preserved in sedimentary rocks deposited between ~2.8 to 2.5 billion years ago reveal unprecedented levels of ^12^C enrichment that are most parsimoniously attributed to the incorporation of biomass formed by methanotrophs (Hayes, 1994; Eigenbrode and Freeman, 2006). Under aerobic conditions, CH_4_ is oxidized with molecular O_2_ to form biomass through the serine or ribulose monophosphate pathways, and CO_2_ (Chistoserdova et al., 2005), where the minimum concentrations of CH_4_ and O_2_ required to drive the microbial process are estimated at 5-10 μM and 12-20 μM, respectively (Harris and Hanson, 1980; Lidstrom and Somers, 1984). Under anaerobic conditions NO_3^-^_, NO_2^-^_, SO_4_^2-^, Fe^3+^, or Mn^2+^ may also be used to oxidize CH_4_ to form simple organic compounds, like acetate (Yan et al., 2018). If photosynthetically derived resources were high — as expected by the pervasive occurrence of thick organic-rich black shales and carbonates in Neoarchean sedimentary successions worldwide (e.g. Zhelezinskaia et al., 2014) — then almost any local increase in O_2_ could have initiated overflow metabolism in extant heterotrophic bacteria. Through microbial community dynamics, this could have resulted in the enhanced release of CH_4_ to the atmosphere, as described in the model above. Distributed globally, oceanic O_2_ oases could then have been the source of Neoarchean CH_4_ hazes.

Independent evidence for the presence of repetitious Neoarchean CH_4_ haze events comes from multiple sulfur isotope measurements of pyrite from the same samples where organic matter was isolated and analyzed. In Neoarchean marginal marine sedimentary successions on multiple continents, carbon isotope evidence links enhanced methanotrophic biomass inputs in organic-rich sedimentary rocks to the presence of a CH_4_ haze that would have attenuated the UV flux and isotopically altered the products of atmospheric SO_2_ photochemistry (see Domagal-Goldman et al., 2008), predominantly sulfate aerosols and elemental sulfur (Kasting, 2001). The photochemical reactions resulted in mass-independent fractionation, or redistribution, of the sulfur isotope masses (S-MIF) in both of the products that rained down and accumulated in surface environments. Notable deviations in the ratio of multiple rare sulfur isotope abundances (Δ^33^S/δ^34^S and Δ^36^S/Δ^33^S) (see **Methods**) relative to normal distribution of Archean S-MIF data (typically referred to as Archean Reference Arrays, or ARAs) coincide in several discrete stratigraphic intervals with negative excursions in carbon isotope values suggestive of methanotrophic inputs of organic carbon (Zerkle et al., 2012; Farquhar et al., 2013; Izon et al., 2015; 2017, Williford et at., 2016). The most recent of these studies (Izon et al., 2017) used available radiometric constraints to estimate the duration of a single haze event at ~1 million years. Through accelerated H_2_ loss to space, the haze events are postulated by these authors to have hastened planetary oxidation. Atmospheric models of Neoarchean hazes suggest that CH_4_ concentrations may have been at the level of 20-30 ppm (e.g. Zerkle et al., 2012), as opposed to ~2 ppm in the modern atmosphere.

Photochemical experiments with SO_2_ have shown that isotope effects and resulting sulfur isotope fractionations depend on both reaction condition and the wavelength of incident radiation. Recent results (Whitehill et al. 2012; Endo et al., 2016) show that shallow positive slopes for Δ^33^S/δ^34^S (<0.1) are produced by photolysis reactions at wavelengths shorter than ~220 nm, and steeper slopes (>3) are produced when longer wavelengths (240 – 340 nm) cause photoexcitation. Similarly, photolysis of SO_2_ yields steeper negative Δ^36^S/Δ^33^S (<-2), while photoexcitation reactions yield positive Δ^36^S/Δ^33^S values (e.g. Whitehill et al., 2012). Applying these observations to the Neoarchean and early Paleoproterozoic atmosphere, shifts to shallower Δ^33^S/δ^34^S and steeper Δ^36^S/Δ^33^S slopes would be anticipated with the transmission of shorter-wavelength UV light resulting in greater proportional photolysis reactions. This may result from the shielding of UV wavelengths by a photochemical CH_4_ haze (e.g. Wolf and Toon, 2010), although quantitative details are lacking.

Pertinent to our model, it appears possible that anoxygenic photosynthesis may have been dominant during intervals when a CH_4_ haze blanketed the planet. The presence of a methane haze would have shielded surface environments from harmful UV radiation that might then allow ammonia (a strong greenhouse gas) to accumulate in the lower atmosphere, and thus have shifted upward the peak wavelength of solar radiation reaching the oceans (Sagan and Chyba, 1996; Wolf and Toon, 2010). Insofar as a Neoarchean CH_4_ haze would scatter and absorb UV and visible wavelengths (Wolf and Toon, 2010), it may have influenced photosynthetic reactions and primary productivity. In this regard, the pigments used during anoxygenic photosynthesis differ from those associated with oxygenic photosynthesis in their molecular details and peak wavelength of light absorbed. Notably the peak absorption wavelength for chlorophyll-a used by oxygen-producing plants and cyanobacteria is 100-200 nm less than the peak absorption wavelength used by bacteriochlorophylls during anoxygenic photosynthesis. The development of a significant methane haze may thus have shifted primary production of organic matter towards anoxygenic photosynthesis where the electrons necessary to drive the metabolic activity ultimately come from the oxidation of H_2_S or Fe (II) (rather than from H_2_O in oxygenic photosynthesis). If correct, the dominance of the anaerobic metabolism over oxygenic photosynthesis would have decoupled inputs of R and O_2_, matching a key assumption of our model.

While our resource-driven ecological model is neither calibrated for amount nor time, the consistent, robust periodic dynamics observed in the system under some parameter regimes nonetheless accord with the episodic release of CH_4_ from surface environments assuming a constant, but not overwhelming, source of O_2_. According to our ecological model, if this oxidant was globally distributed at low concentrations, and if resources were high, a CH_4_ haze would likely result. Izon et al. (2017) suggest that the coupled carbon/sulfur isotope anomalies were associated with biospheric responses to increased nutrients, with the CH_4_ flux controlled by the availability of organic carbon and sulfate. Increased loading of the oceans with nutrients (e.g., nitrogen, phosphorus) is most often associated with uplift, mountain building and weathering of exposed continental masses, all of which could erode, disrupt, or accelerate sedimentation on marginal marine platforms. In the relatively continuous sedimentary successions where these episodic coupled anomalies have been recorded, there is no physical evidence for any temporal sedimentary disruption in the form of unconformities or major hiatal surfaces. As an alternative hypothesis, we propose that if nutrients (including ferrous iron, which would have been soluble in the ocean under anoxic conditions) were widely available in the Neoarchean ocean, then the dynamic release of CH_4_ and the episodic development of CH_4_ hazes in the atmosphere could simply have been the result of constant low level (or gradually increasing) inputs of photosynthetic O_2_ to anaerobic environments where overflow metabolism was dominant.

Geochemically documented CH_4_ haze events are currently known only from the Neoarchean, but the episodic and potentially globe-encompassing ice ages of the succeeding early Paleoproterozoic Era may similarly be understood in terms of fluctuations of O_2_ and greenhouse gases in the ancient atmosphere (Pavlov et al, 2000; Bekker et al., 2005; Kopp et al., 2005; Goldblatt et al., 2006; Claire et al., 2006; Bekker and Kaufman, 2007). Insofar as the early Paleoproterozoic glacial epoch is related to the transition from the anoxic CH_4_-rich atmosphere to an oxic CO_2_-rich atmosphere, the multiplicity of ice ages (including at least three discrete events) in this interval is arguably related to oscillations in O_2_ and CH_4_ production and utilization. Related to the dynamic ecological model presented here, the rise of photosynthetic O_2_ prior to each of these Paleoproterozoic glaciations would have progressively drawn down atmospheric CH_4_ levels resulting in surface refrigeration. Primary productivity would then be severely limited beneath ice-covered oceans, thereby constraining the production of both fermentative products (FP) and O_2_. Atmospheric CO_2_ levels during the glaciations likely rose due to unabated volcanic fluxes and limited chemical weathering of exposed continental rocks (e.g., Hoffman et al., 1998). However, once threshold concentrations of CO_2_ were reached, the glaciations would have ended rapidly, resulting in high rates of silicate weathering on land, and the enhanced delivery of sediments, oxidants (including sulfate and nitrate), nutrients, and alkalinity to the oceans. Melting glaciers and sea ice coupled with higher post-glacial temperatures would have flooded continental margins and stimulated both oxygenic photosynthesis and methanogenesis.

Support for such an environmental scenario comes from carbon isotope analyses of drill core samples collected from across the oldest recorded ice age deposit (the Ramsay Lake diamictite) in southern Ontario, which reveal strongly negative δ^13^C values (with a nadir of ca. −41 ‰) in organic matter from sediments that accumulated in the glacial aftermath (Bekker and Kaufman, 2007). The magnitude and duration of ^13^C depletion in the post-glacial Pecors Formation (expressed over 100 meters of sedimentary strata) is consistent with long-term methanogenic production and methanotrophic oxidation of CH_4_ in a shallow oceanic setting. Multiple rare sulfur isotope abundances in pyrite extracted from the same Pecors samples reveal even more profound negative deviations in Δ^36^S/Δ^33^S values relative to the ARA (Wing et al., 2002; 2004), which may similarly record atmospheric processes during this (and other) post-glacial early Paleoproterozoic CH_4_ haze events. Considering our model results, we suggest that the end of the early Paleoproterozoic glacial epoch around 2.2 billion years ago likely marked the end of extended methane-dominated atmospheres of the early Earth. This repetitive series of ice ages was followed by a remarkably long-lived (>100 million years) interval of highly positive carbon isotope compositions in carbonates known as the Lomagundi-Jatuli event (Karhu and Holland, 1996; Bachan and Kump, 2015; Mänd et al., 2020). Insofar as significant ^13^C enrichments in marine carbonates are driven by the burial and sequestration of ^12^C-rich organic matter (Broecker, 1970; Hayes, 1983), the attendant photosynthetic production and release of O_2_ at this time would most likely have marked the transition to an oxygenated atmosphere with carbon dioxide acting as the major greenhouse gas thereafter.

Episodically enhanced CH_4_ release and buildup in the atmosphere later in Earth history may be related to the tectonic reorganization of the mobile continents, which modulate the delivery of reduced gases from spreading ridges and weathering fluxes from uplifted mountains. These variable fluxes would have impacted the photosynthetic production of organic matter and degree of ocean oxygenation during discrete climatic and biological events. Insofar as our model results indicate that these factors control CH_4_ fluxes to the atmosphere, we speculate that younger ice ages of the Neoproterozoic and Paleozoic eras, as well as some Phanerozoic mass extinctions, may be linked to the dynamic microbial processes modeled here.

## Conclusion

Patterns of CH_4_ production and release in both driven and non-driven versions of a novel microbial ecological model, based upon the metabolic repercussions of the Warburg Effect, suggest that resources and O_2_ play important roles in defining the dynamics of microbial ecosystems over both short and long-time scales. We demonstrate that forced low level inputs of O_2_ result in pulsed releases of CH_4_ via overflow metabolism, a pattern that may explain periodic CH_4_ haze events and episodic glaciation deep in Earth’s history. Such pulsed methane releases also occur in modern-day wetlands where tidal flooding repeatedly tips the balance between oxic and anoxic conditions (see Zhu et al., 2017), suggesting similar mechanisms may be at work. The qualitative behavior of this ecosystem model provides a platform for understanding the dynamics of mixed microbial communities in ancient, modern, and future ecosystems. More detailed models that blend empirical gas-release data with calibrated input parameters should shed further light on the links between microbial communities and the timing of atmospheric dynamics.

## Methods

### Modeling Microbial System Dynamics

We model the dynamics of the interacting microbial system as a system of ordinary differential equations (ODEs). First, we define functions pertaining to the Warburg Effect (eq. 1–2) and then describe the equations for each of the microbial functional groups.

We begin by defining two functions that describe Warburg Effect dynamics in aerobic heterotrophic bacteria (A):

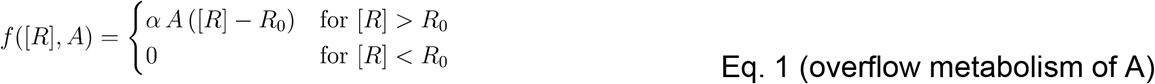

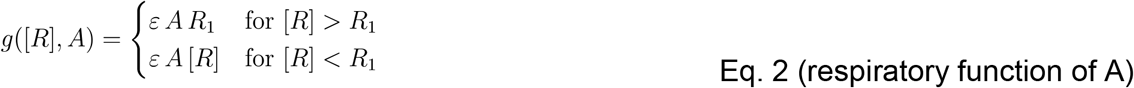

The first function *f*(*R, A*) denotes the process of overflow metabolism, where, despite the presence of ample oxygen, heterotrophic aerobic bacteria (A) produce fermentation products when resources are abundant. In this function, at resource concentrations higher than base levels (represented by *R_0_*), there is a linear increase in overflow products (see Basan et al., 2015, Swain and Fagan, 2018). The second function *g*(*R, A*) denotes the respiratory function (see Eq. 3 below) for these organisms (with the maximal resource usage condition represented by resource concentration of *R_1_*). Both *α* and *ϵ* are proportionality constants that denote the efficiency of overflow metabolism and respiration, respectively, and fall in the range [0, 1].

The differential equations below pertain to changes through time of the dominance of metabolic functional groups as the different microbial functional groups (A, B, C, F, and G as defined in Fig. 1) compete for available substrates under variable environmental conditions.

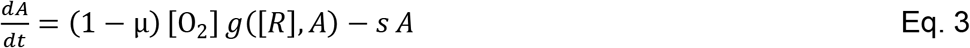

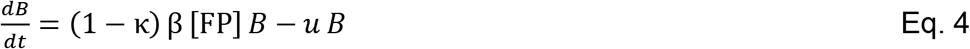

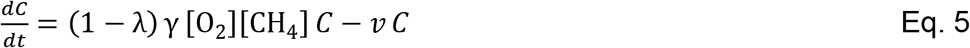

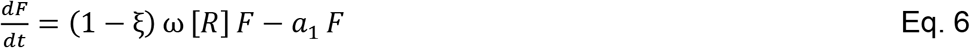

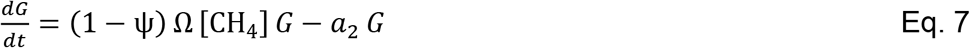

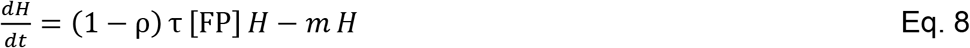

The first term on the right-hand side (RHS) of each differential equation is positive and denotes the process of assimilation of substrates needed for growth (increase in biomass) and energy production for the metabolic class. The second RHS term in each equation is negative and denotes the energy requirement for homeostasis (see Table 1). In equations 3–8, the terms governing resource incorporation efficiency, generically denoted : (1 − *X*) with X replaced by the consortium-specific parameters (*μ, κ, λ, ξ* and *ψ*), specify the proportion of the required substrate in the extracellular environment that is incorporated into cells, which can then be used for making biomass by those microbes. Variation in these efficiencies is assumed to be modulated by biochemical and physical constraints of the different microbial functional groups. The remaining reservoir of resources outside of the cell either leave the system as gases or are available for later incorporation. All twelve coefficients of efficiency (*μ, κ, λ, ξ, ψ, ρ, ε, β, γ, ω, Ω and τ*) lie in the range [0, 1]. The coefficients related to homeostasis (*s, u, v, a*_1_, *a*_2_, *or m*) represent the basal metabolic requirement values for microbial functional groups A, B, C, F, G, and H respectively. All parameters and their interpretations are summarized in Table 1.

**Table 1:** Differential equation parameters governing the dynamics of a system of five microbial functional groups. Parameters summarize the efficiency of resource incorporation, the efficiency of resource utilization, and the energy requirements for homeostasis.

| Functional group | Resource<br>incorporation<br>efficiency | Resource<br>utilization<br>efficiency | Energy<br>requirements<br>for<br>homeostasis |
| --- | --- | --- | --- |
| A (aerobic heterotrophs) | $(1 - \mu)$ | $\varepsilon$ (for R) | $s$ |
| B (methanogens) | $(1 - \kappa)$ | $\beta$ (for FP) | $u$ |
| C (aerobic<br>methanotrophs) | $(1 - \lambda)$ | $\gamma$ (for O<sub>2</sub> and<br>CH<sub>4</sub>) | $v$ |
| F (fermenters) | $(1 - \xi)$ | $\omega$ (for R) | $a_1$ |
| G (anaerobic<br>methanotrophs) | $(1 - \psi)$ | $\Omega$ (for CH<sub>4</sub>) | $a_2$ |
| H (anaerobic<br>heterotrophs) | $(1 - \rho)$ | $\tau$ (for FP) | $m$ |

The following set of differential equations reflect dynamic changes in the abundance of resources or metabolic products in the five microbial functional groups through time:

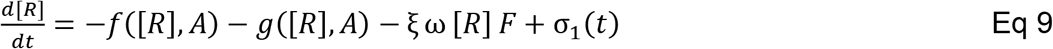

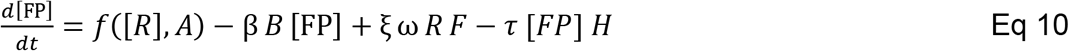

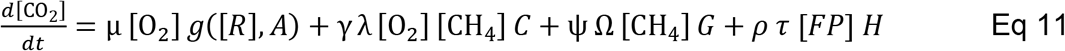

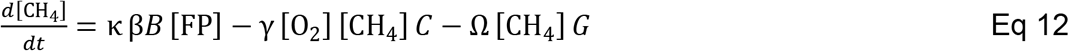

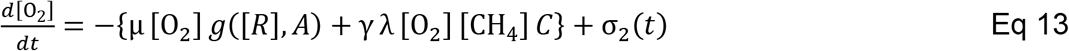

In equations 8–12 the RHS positive terms indicate either input or production of the resources, while negative terms indicate metabolic resource consumption. All the terms involve interaction of the resources with one or more of the microbial functional groups, which are modelled in equations 3–8, except for the ‘forcing terms’ *σ*_1_(*t*) *and σ*_2_(*t*). These terms (see Eqs. 9 and 13) denote the time-dependent input rates of R and O_2_ that are controlled externally. In this model, CO_2_ is considered a sink that evades to the atmosphere and is decoupled from the autotrophic processes. This assumption makes R an independent variable, and further ensures that substrate production is not photoautotrophic (the case in many scenarios). Moreover, as developed here, the ecological model is not spatially compartmented in order to make the results more general in nature.

There exist many methanogenic systems where oxic and anoxic parts are heterogeneously mixed, albeit at generally small scales or temporally, such as in wetlands (Kirsche et al., 2013). In this ecological model we have simplified the process of methanogenesis in several ways: i) FPs can be one of many substrates, including formate, acetate, and lactate, all of which can be used for methane production by various methanogens; and ii) for hydrogen (H_2_) + CO_2_ methane production, we assume that reduced reactants are unlimited, and insofar as CO_2_ is a sink and evades the system as a gas, it is unlimited as well (i.e., it is not used for autotrophic processes that deliver new resources to the community). For the anaerobic oxidation of methane by functional group G, we lump all of the potential anaerobic methanotrophs here, but recognizing that both CO_2_ and carbonate alkalinity may be formed, we include a conversion efficiency term (*Ω*) to calculate the relative concentrations of CO_2_ alone.

This coupled differential equation system was solved through Wolfram Mathematica v. 12.1 and R v. 3.6.2. A regular parameter sweep was first performed for the non-driven case (no external input: *σ*_1_, *σ*_2_ = 0) using Latin Hypercube sampling (LHS) for 500,000 different scenarios using R packages *deSolve* (Soetaert et al., 2010) and *lhs* (Carnell, 2020). We then calculated the partial rank correlation coefficients (PRCCs) to explore the effect of each parameter and initial condition on the concentrations of gases (CH_4_, O_2_ and CO_2_) through R package *sensitivity* (Iooss et al., 2020).

To further explore the trends in these gas emissions, we calculated the relative proportions of these gases under a set of base parameters that depict biologically possible scenarios (according to data from Payne and Wiebe, 1978; Geyer et al., 2016; Bowles et al., 2019; and Morris and Schmidt, 2013). All the efficiencies were set at half maximum for the base case. We then varied one parameter at a time to look at the dependence on that specific variable on gas emissions as a function of resource and oxygen initial conditions. In doing so, we collated the data for different initial conditions (such as different initial ratios of the microbial functional groups) and plotted the averaged values of the equilibrium state as a function of initial resource and oxygen inputs. For driven dynamics, we followed the same procedure of parameter dependence, but now including the two R and O_2_ external input parameters, *σ*_1_ and *σ*_2_. To calculate the period and amplitude of the driven dynamics, values were numerically estimated based on dynamic results from the first few oscillations. Periodicity statistics for the dynamics resulting from both high and low extreme values of *σ*_2_ are estimates given the difficulty of estimating them for very long or extremely short oscillations that occur at these values of *σ*_2_.

For looking at the sensitivity of the model to addition or removal of microbial functional groups, we ran the same tests for a model without H (anaerobic heterotrophs). To measure sensitivity of the demarcation of the oscillatory phase in the space of resource and oxygen addition (*σ*_1_ − *σ*_2_ *space*), across a wide variety of parameter combinations, we first used LHS to sample 100,000 combinations through the parameters space. We calculated, for each set of parameters, twenty points on the *σ*_1_ − *σ*_2_ space, estimated the relative value for which there is a phase change from oscillatory behavior to no oscillations, and from these values calculated the slope of the line which demarcates the space for each set of LHS parameters. We performed PRCC analysis to find the dependence of the slope value on the parameters.

### Notation for Sulfur Isotope Compositions

Sulfur-isotope ratios are conventionally reported in delta (δ) notation and reflect the permil (‰) deviation of the ratio of the least abundant isotope (^33^S, ^34^S, or ^36^S) to the most abundant isotope (^32^S), relative to the same ratio in an international reference standard (Vienna Canyon Diablo Troilite, V-CDT). For example, δ^34^S = [(^34^S/^32^S)_sample_/(^34^S/^32^S)_V-CDT_ − 1] x 1000. The majority of terrestrial processes fractionate S-isotopes mass-dependently, whereby δ^33^S ≈ 0.515 * δ^34^S and δ^36^S ≈ 1.91 * δ^34^S. Departure from mass-dependent behavior, or mass-independent fractionation (S-MIF), is expressed in capital-delta (Δ) notation as either non-zero Δ^33^S = [(^33^S/^32^S)_sample_/(^33^S/^32^S)_V-CDT_ − (^34^S/^32^S)_sample_/(^34^S/^32^S)_V-CDT_]^0.515^ or Δ^36^S = [(^36^S/^32^S)_sample_/(^36^S/^32^S)_V-CDT_ − (^34^S/^32^S)_sample_/(^34^S/^32^S)_V-CDT_]^1.9^.

## Acknowledgements

This research was supported in part by NSF award EAR-1338810 to AJK. AS would like to thank the COMBINE program at the University of Maryland (DGE-1632976) for technical and academic support.

## Code and data availability

All the codes that were used to explore the ODE model and generate the relevant figures are available publicly at the following github repository: https://github.com/anshuman21111/CH4_haze. All extra figures and simulation data, above and exceeding the supplementary materials are also found on this repository.

