## Supplementary figures for "Were Neoarchean atmospheric methane hazes and early Paleoproterozoic glaciations driven by the rise of oxygen in surface environments?"

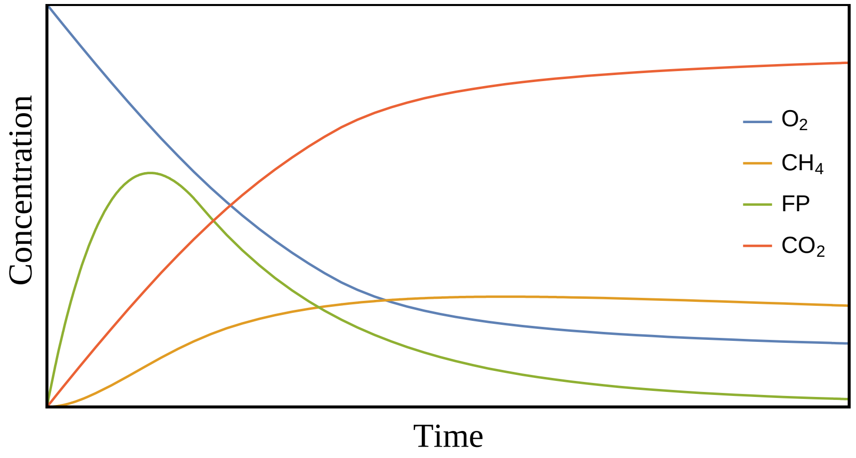


***Figure S1:*** *Dynamic variations in $O_2$, FP, $CH_4$, and $CO_2$ through time with set initial boundary conditions so that (relative) concentrations eventually level out when the system reaches steady-state equilibrium, for a non-driven case. Please note that, here time is in arbitrary units, as pinning an exact time-scale on these would need more information on various modelled microbial parameters, which currently is limited by known literature.*

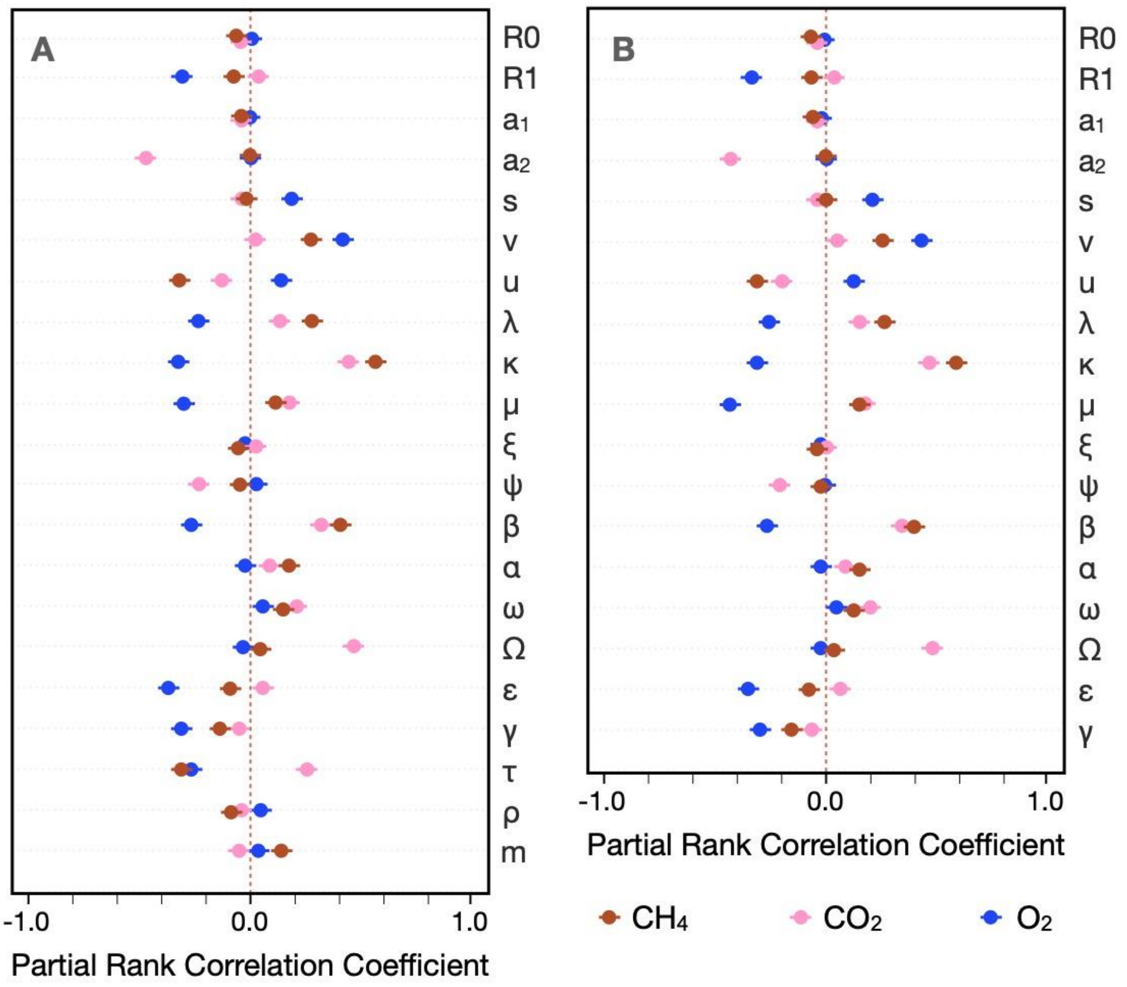


***Figure S2:*** *Latin hypercube sampling (LHS) based Partial rank correlation coefficient (PRCC) (which varies from -1 to 1) of the model parameters with respect to the final gas concentrations in (A) the complete model, (B) in the model with H removed (for checking sensitivity). (A) and (B) have negligible differences, signifying model capacity to perturbations and additions.*

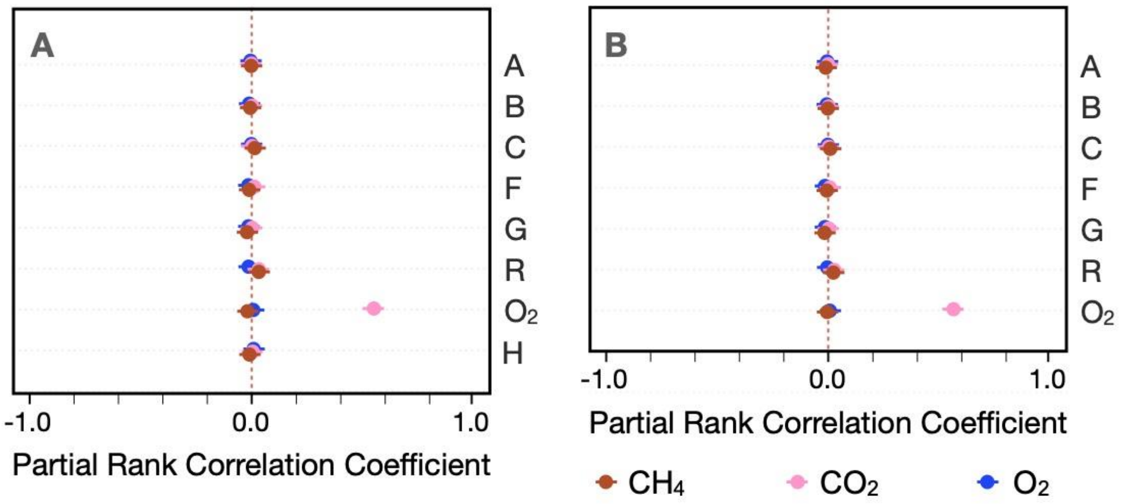


**Figure S3:** Latin hypercube sampling (LHS) based Partial rank correlation coefficient (PRCC) (which varies from -1 to 1) of the model initial conditions with respect to the final gas concentrations in (A) the complete model, (B) in the model with H removed (for checking sensitivity). (A) and (B) have negligible differences, signifying model capacity to perturbations and additions. Please note the extremely small values (absolute value <0.05) of PRCC in all cases except that of $CO_2$ dependence on $O_2$ initial value.

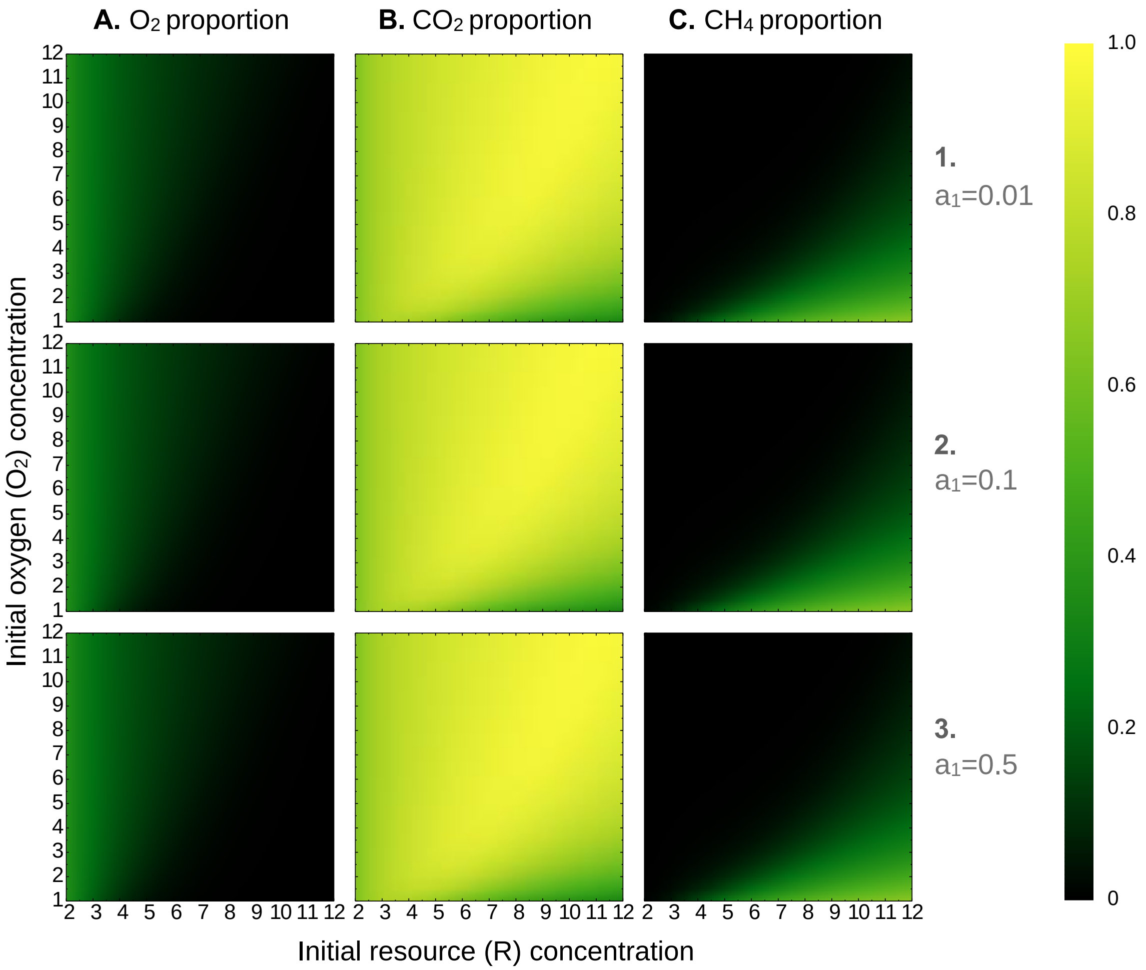


21

22 ***Figure S4A:*** *Variations in $O_2$, $CO_2$, and $CH_4$ proportions (in columns A-C) run to steady state at set $O_2$*
23 *and R input values, with the energy requirement for fermenters ($a_1$) set at 1, 10, and 50% in rows 1-3,*
24 *respectively.*

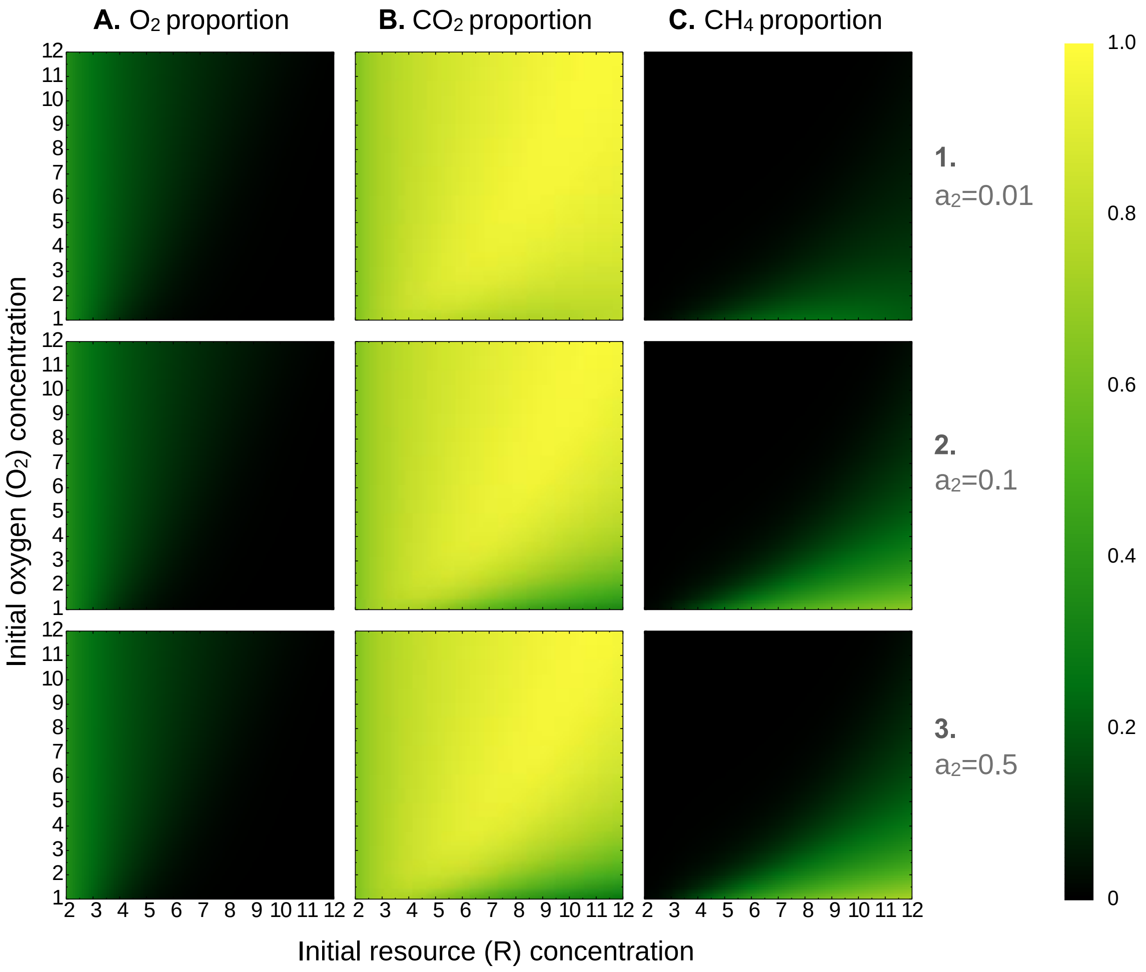


***Figure S4B:*** *Variations in $O_2$, $CO_2$, and $CH_4$ proportions (in columns A-C) run to steady state at set $O_2$ and R input values, with the energy requirement for anaerobic methanotrophs ($a_2$) set at 1, 10, and 50% in rows 1-3, respectively.*

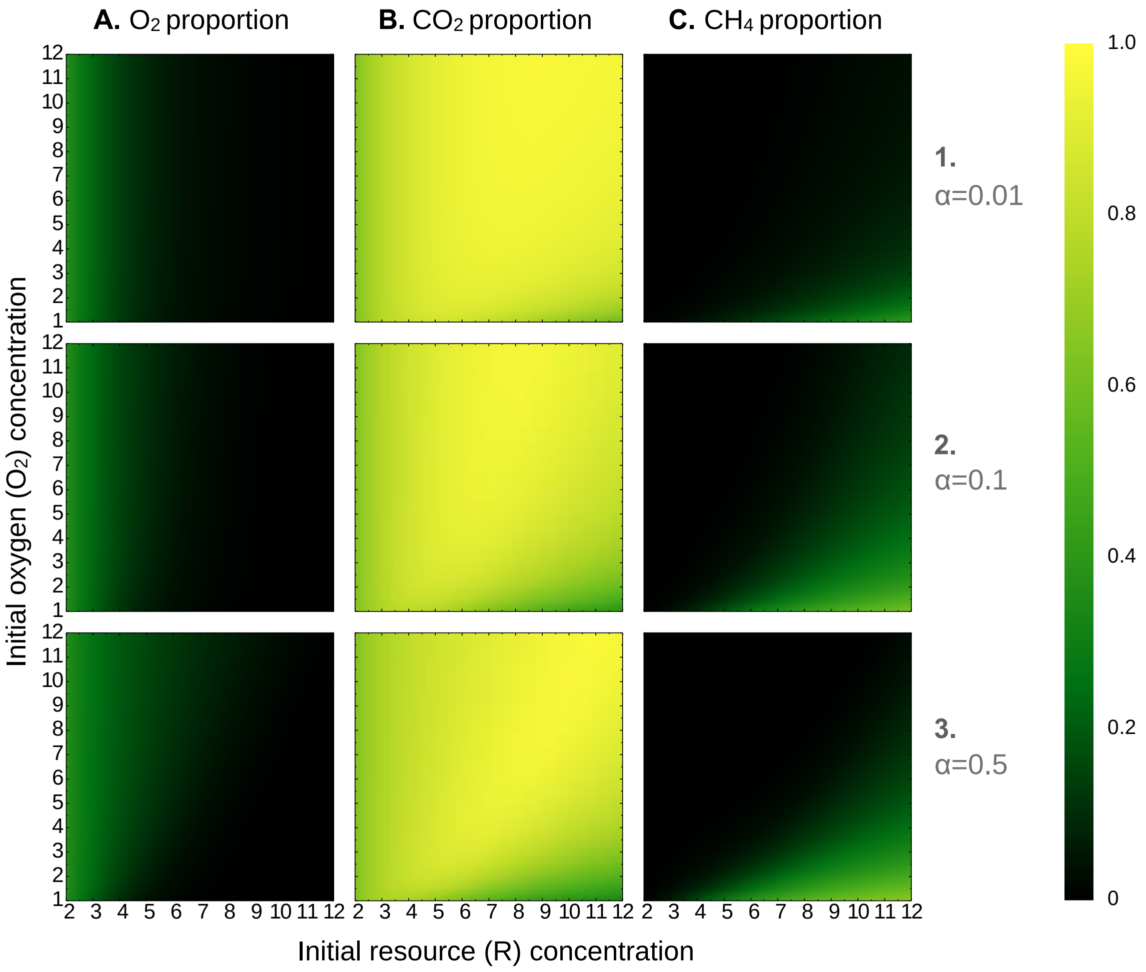


***Figure S4C:*** *Variations in $O_2$, $CO_2$, and $CH_4$ proportions (in columns A-C) run to steady state at set $O_2$ and R input values, with efficiency of overflow metabolism in aerobic heterotrophs (α) set at 1, 10, and 50% in rows 1-3, respectively.*

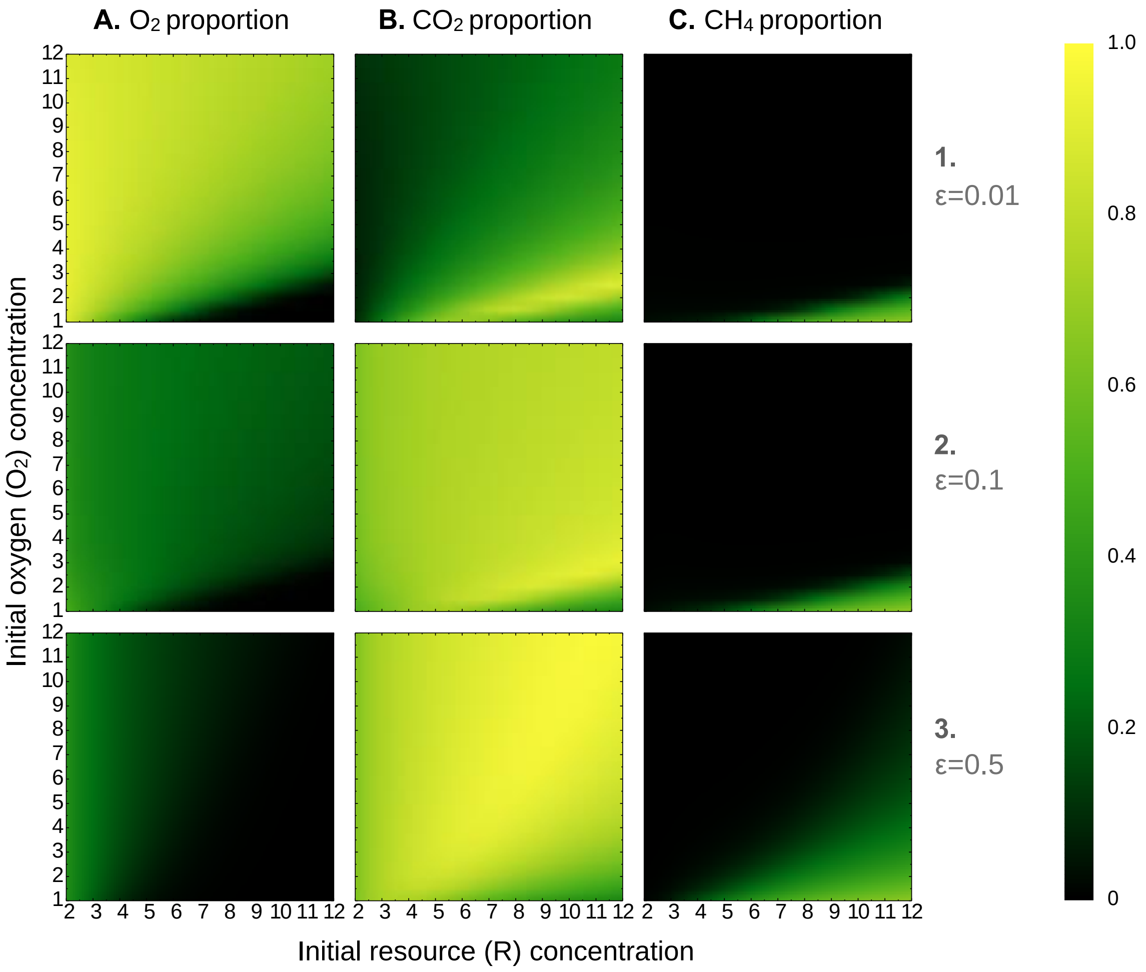


***Figure S4D:*** *Variations in $O_2$, $CO_2$, and $CH_4$ proportions (in columns A-C) run to steady state at set $O_2$ and R input values, with resource utilization efficiency for respiration in aerobic heterotrophs (ε) set at 1, 10, and 50% in rows 1-3, respectively.*

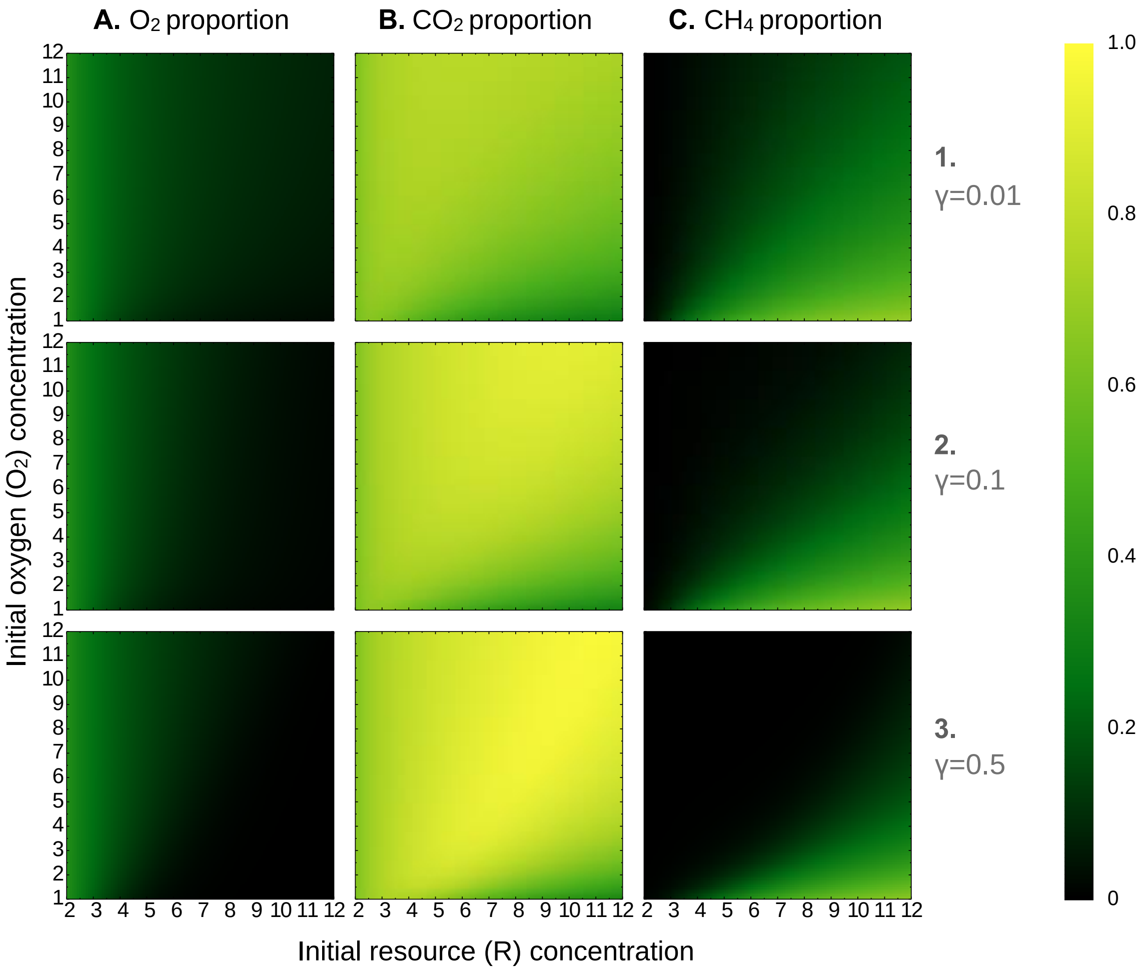


***Figure S4E:*** *Variations in $O_2$, $CO_2$, and $CH_4$ proportions (in columns A-C) run to steady state at set $O_2$ and R input values, with resource utilization efficiency in aerobic methanotrophs (γ) set at 1, 10, and 50% in rows 1-3, respectively.*

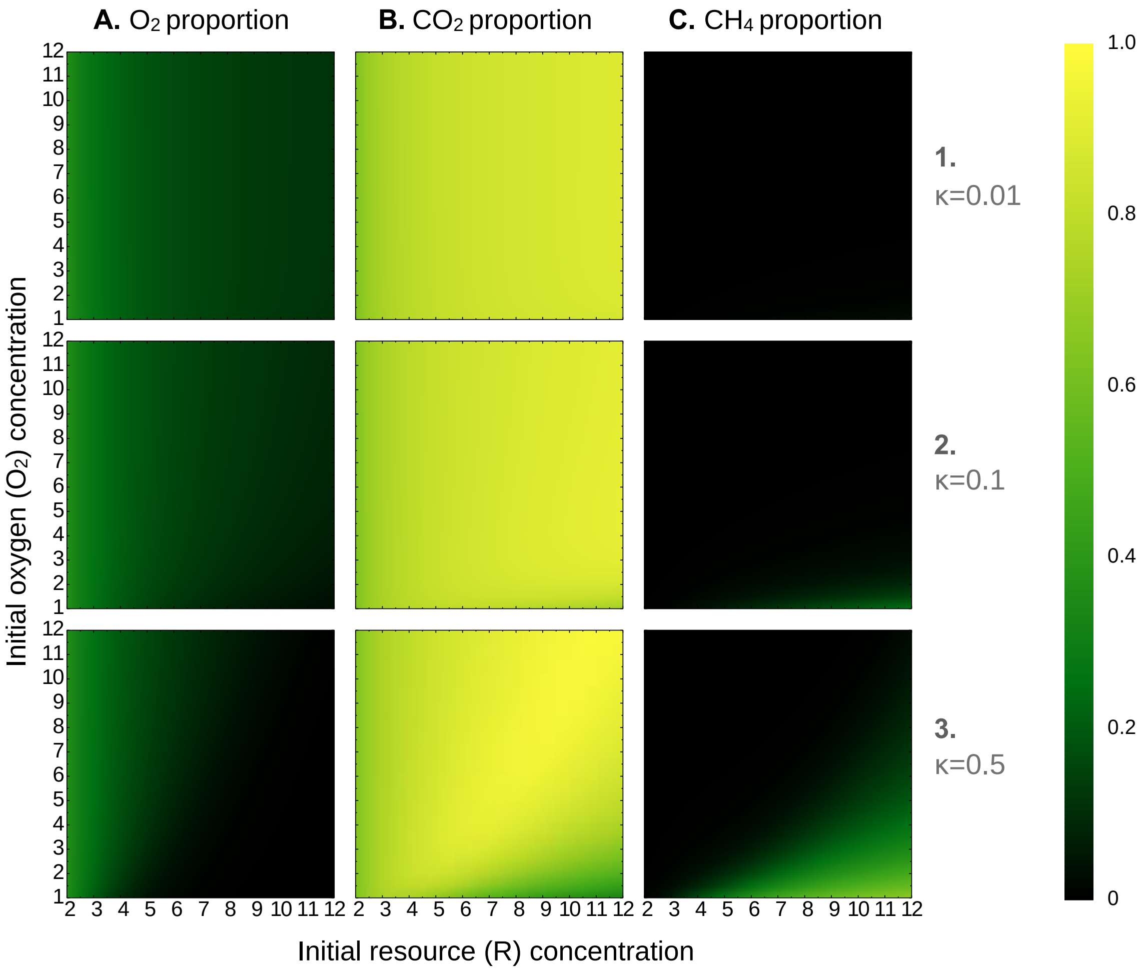


***Figure S4F:*** *Variations in $O_2$, $CO_2$, and $CH_4$ proportions (in columns A-C) run to steady state at set $O_2$ and R input values, with resource incorporation efficiency in methanogens (κ) set at 1, 10, and 50% in rows 1-3, respectively.*

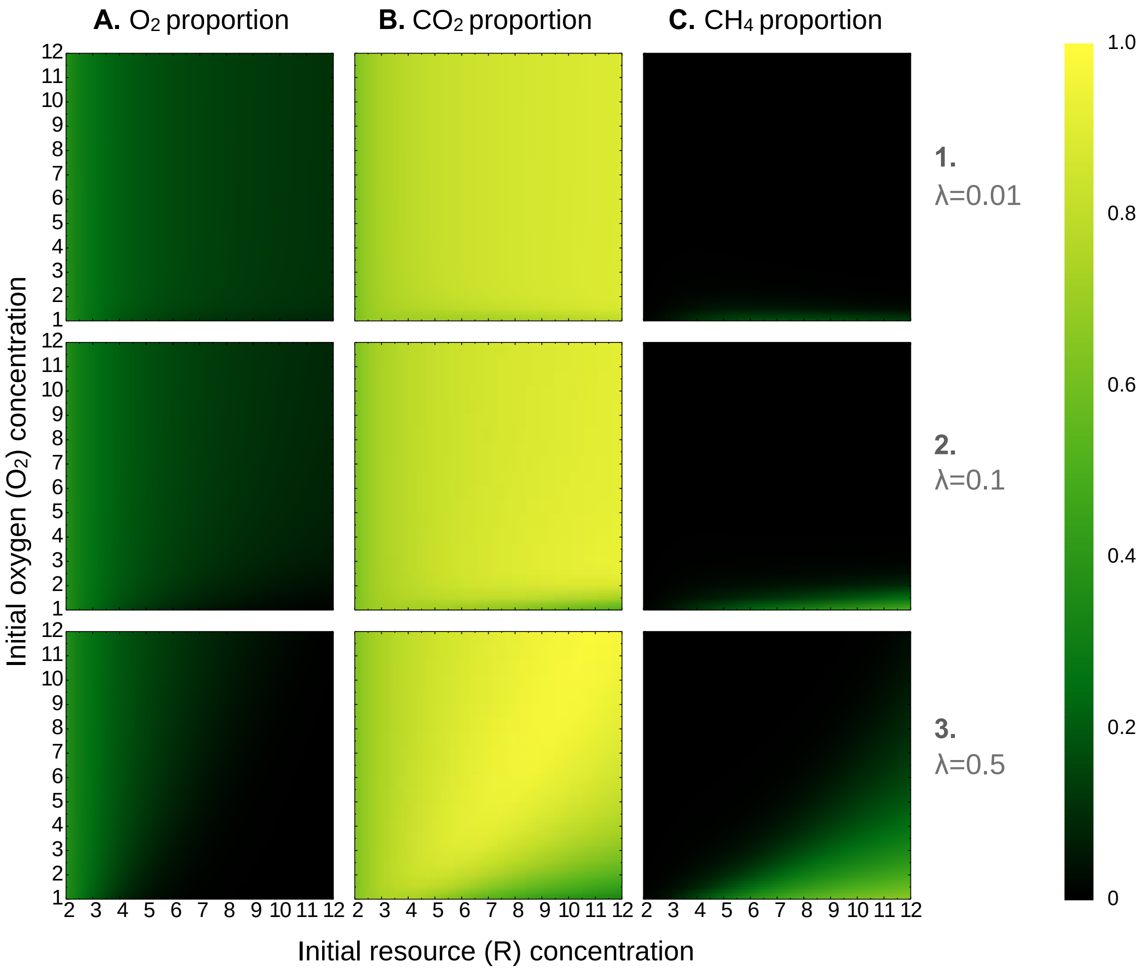


***Figure S4G:*** *Variations in $O_2$, $CO_2$, and $CH_4$ proportions (in columns A-C) run to steady state at set $O_2$ and R input values, with resource incorporation efficiency in aerobic heterotrophs (λ) set at 1, 10, and 50% in rows 1-3, respectively.*

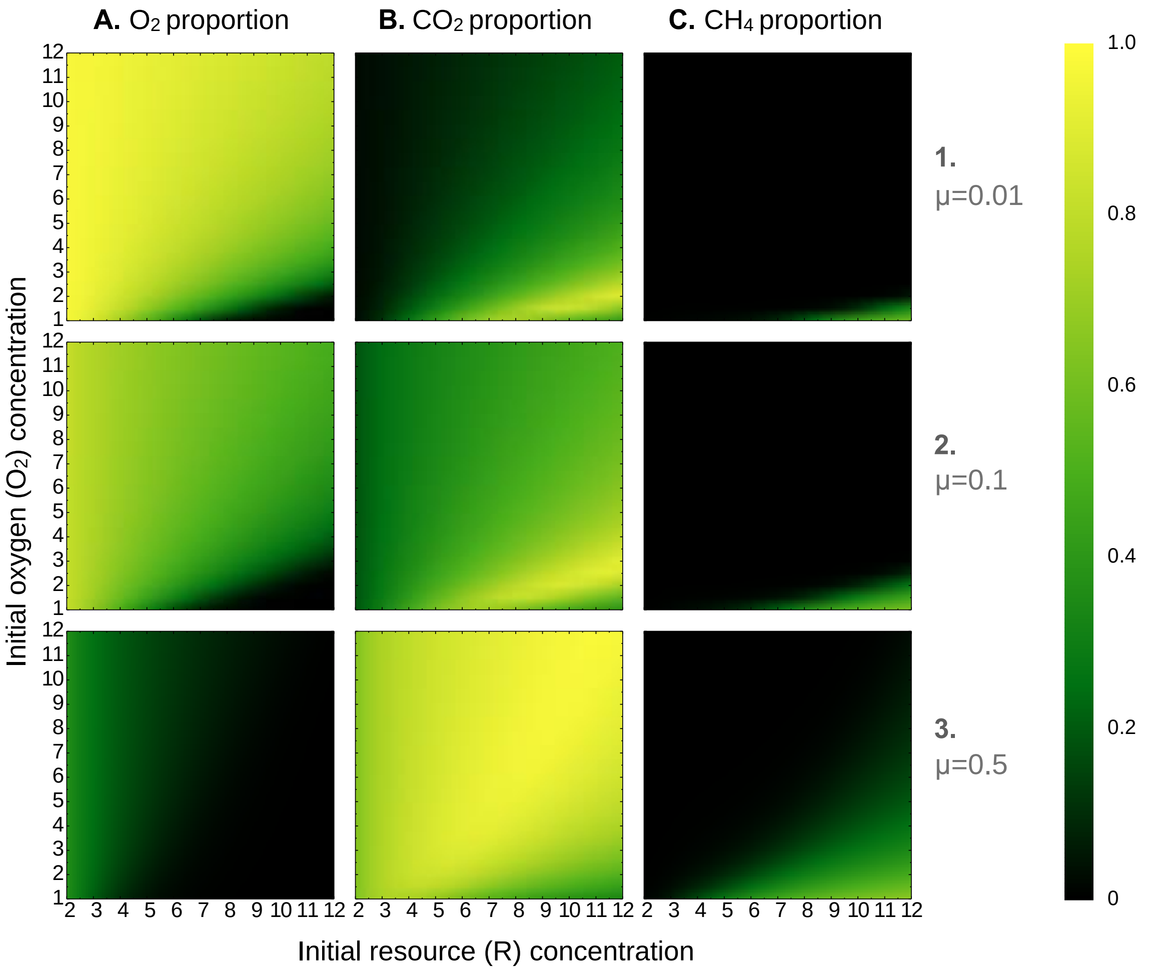


***Figure S4H:*** *Variations in $O_2$, $CO_2$, and $CH_4$ proportions (in columns A-C) run to steady state at set $O_2$ and R input values, with resource incorporation efficiency in aerobic methanotrophs (μ) set at 1, 10, and 50% in rows 1-3, respectively.*

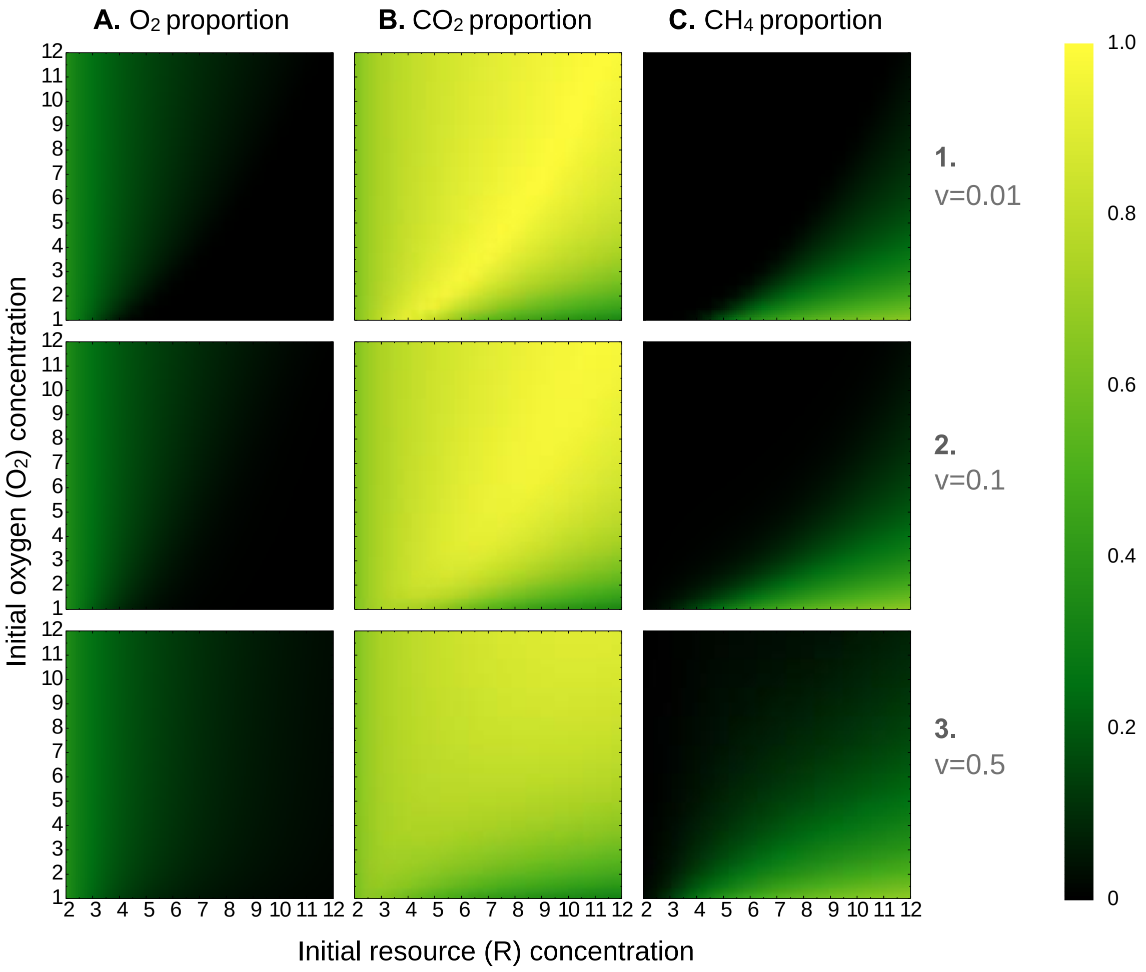


***Figure S4I:*** *Variations in $O_2$, $CO_2$, and $CH_4$ proportions (in columns A-C) run to steady state at set $O_2$ and R input values, with energy requirements of aerobic methanotrophs (v) set at 1, 10, and 50% in rows 1-3, respectively.*

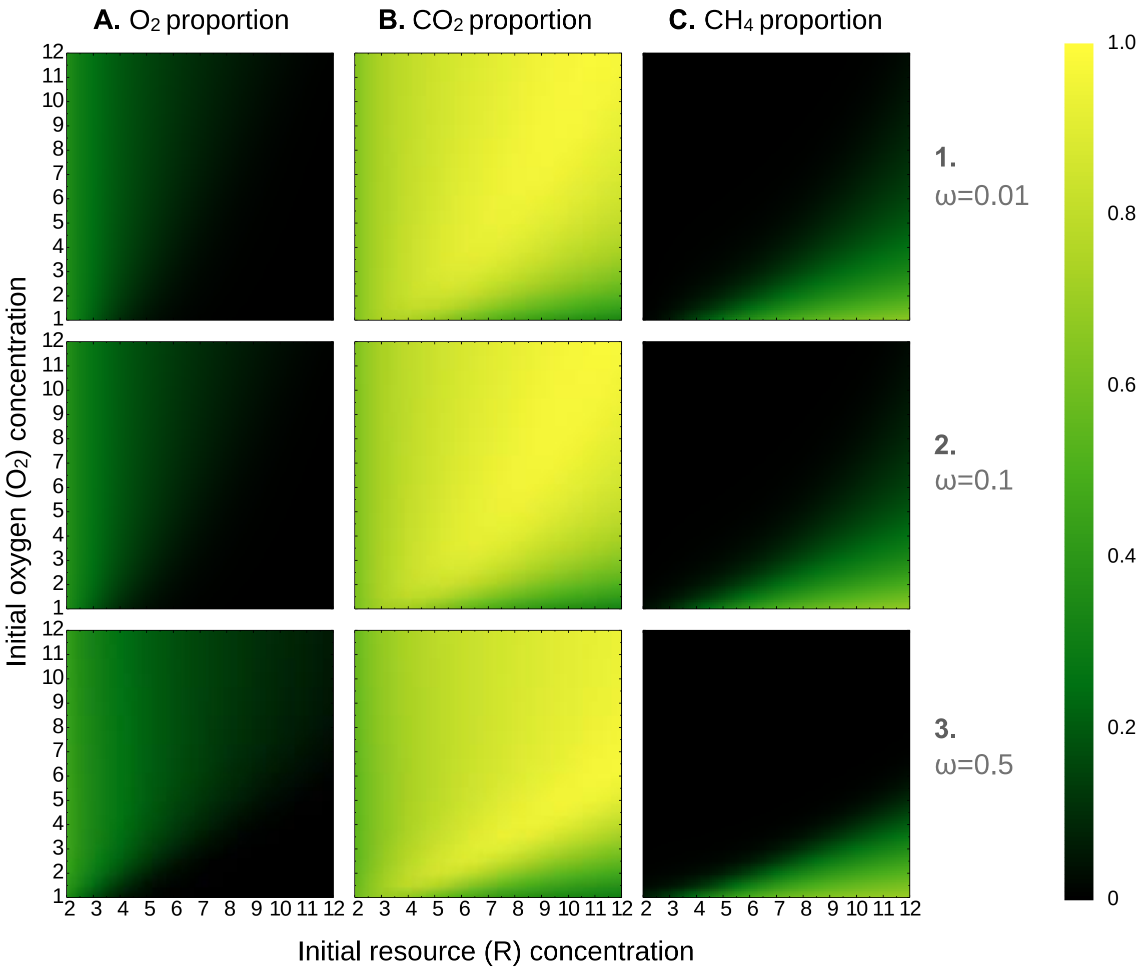


***Figure S4J:*** *Variations in $O_2$, $CO_2$, and $CH_4$ proportions (in columns A-C) run to steady state at set $O_2$ and R input values, with resource utilization efficiency in fermenters (ω) set at 1, 10, and 50% in rows 1-3, respectively.*

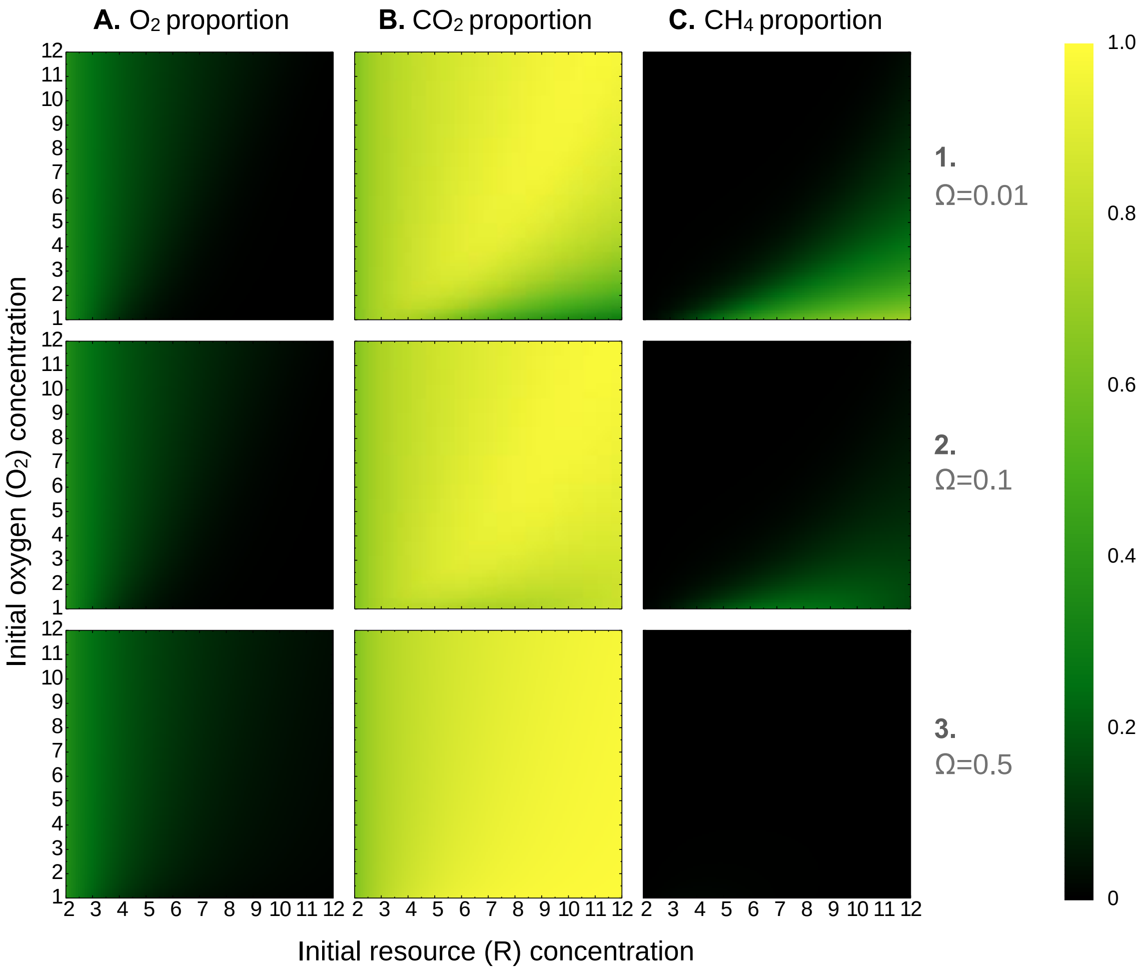


***Figure S4K:*** *Variations in $O_2$, $CO_2$, and $CH_4$ proportions (in columns A-C) run to steady state at set $O_2$ and R input values, with resource utilization efficiency in anaerobic methanotrophs (Ω) set at 1, 10, and 50% in rows 1-3, respectively.*

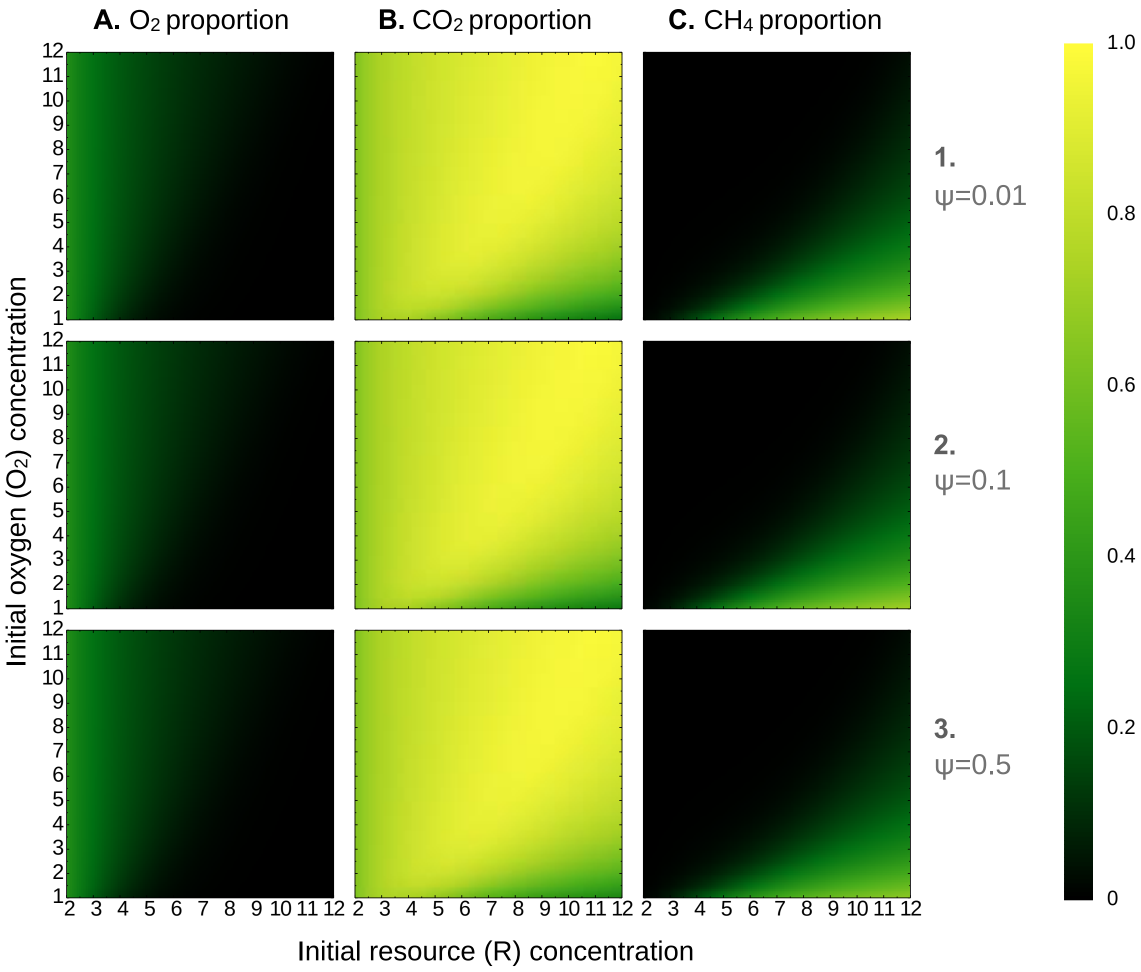


***Figure S4L:*** *Variations in $O_2$, $CO_2$, and $CH_4$ proportions (in columns A-C) run to steady state at set $O_2$ and R input values, with resource incorporation efficiency in anaerobic methanotrophs ($\psi$) set at 1, 10, and 50% in rows 1-3, respectively.*

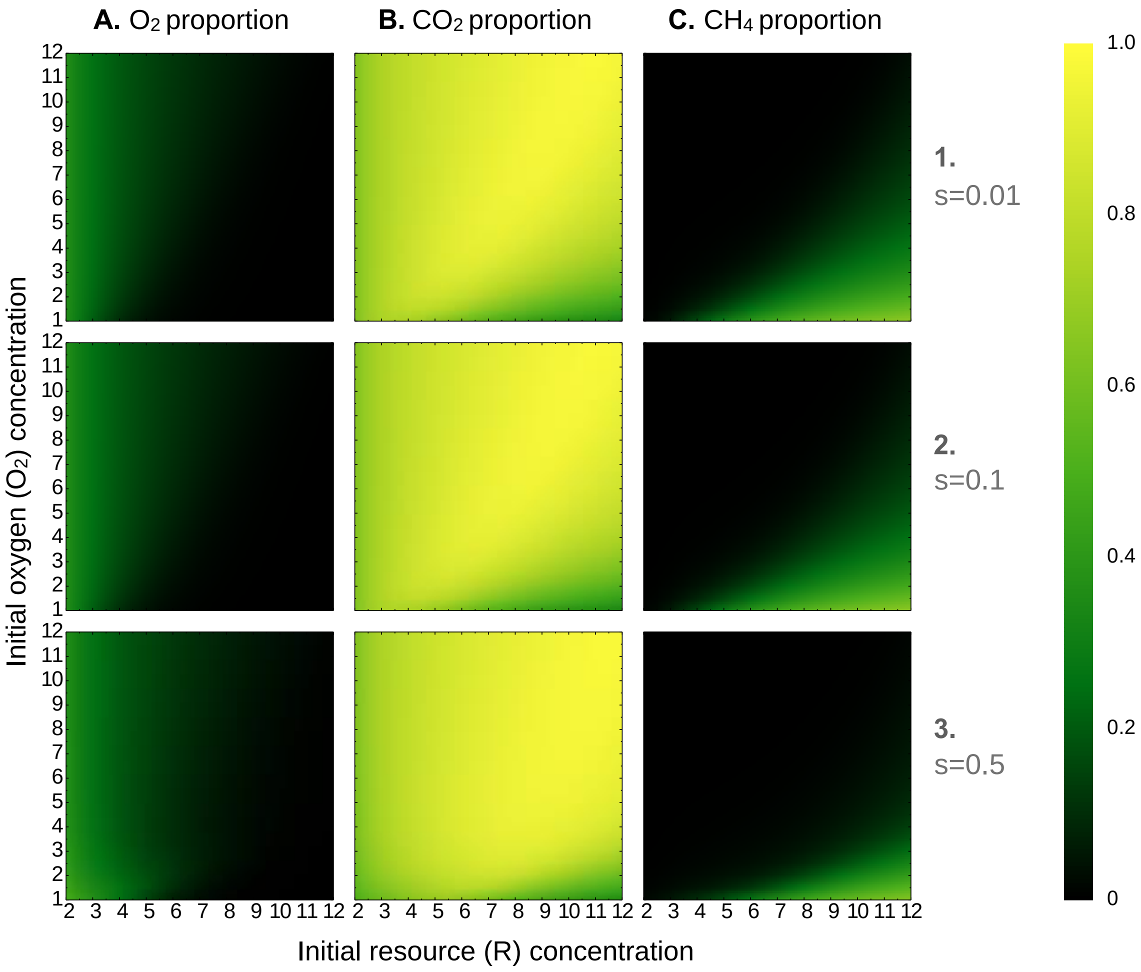


***Figure S4M:*** *Variations in $O_2$, $CO_2$, and $CH_4$ proportions (in columns A-C) run to steady state at set $O_2$ and R input values, with energy requirement for aerobic heterotrophs (s) set at 1, 10, and 50% in rows 1-3, respectively.*

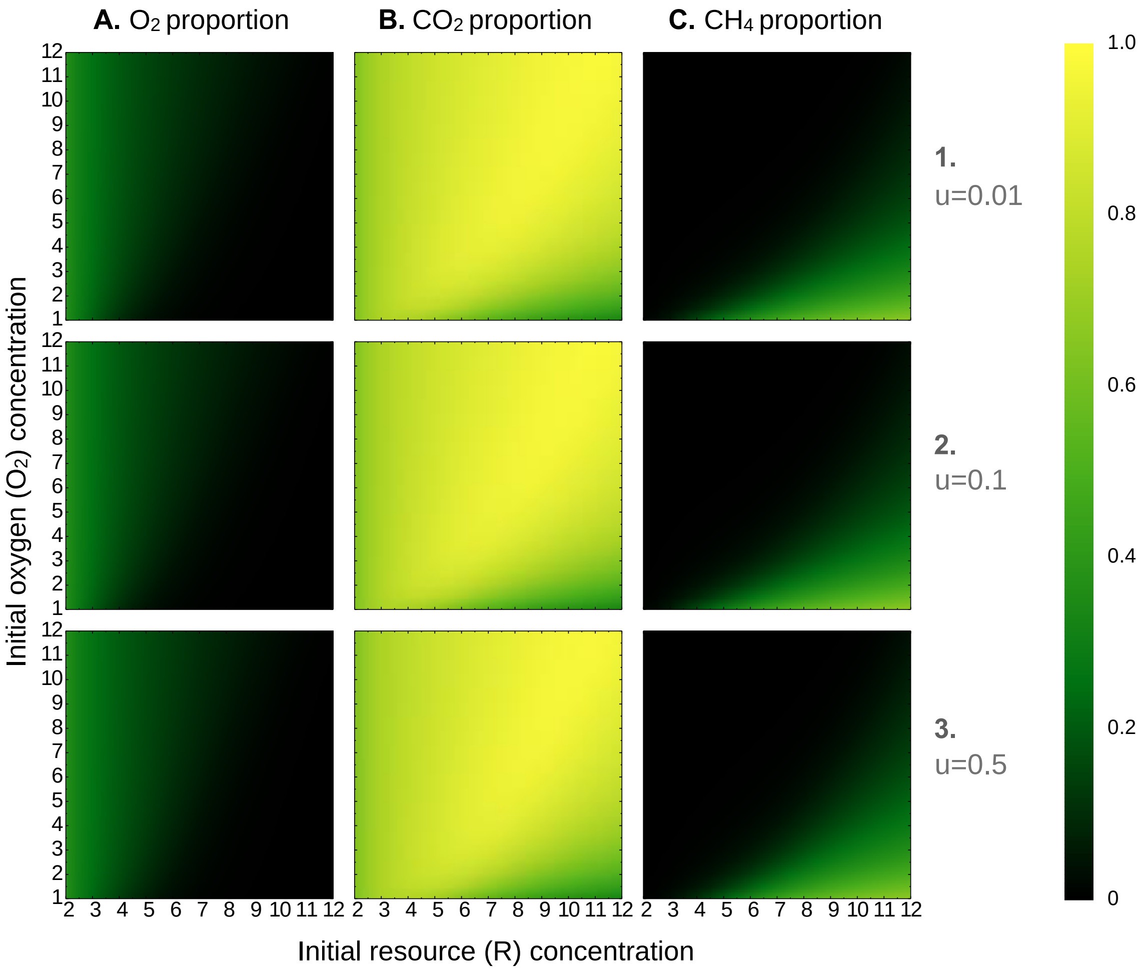


***Figure S4N:*** *Variations in $O_2$, $CO_2$, and $CH_4$ proportions (in columns A-C) run to steady state at set $O_2$ and R input values, with energy requirement for methanogens (u) set at 1, 10, and 50% in rows 1-3, respectively.*

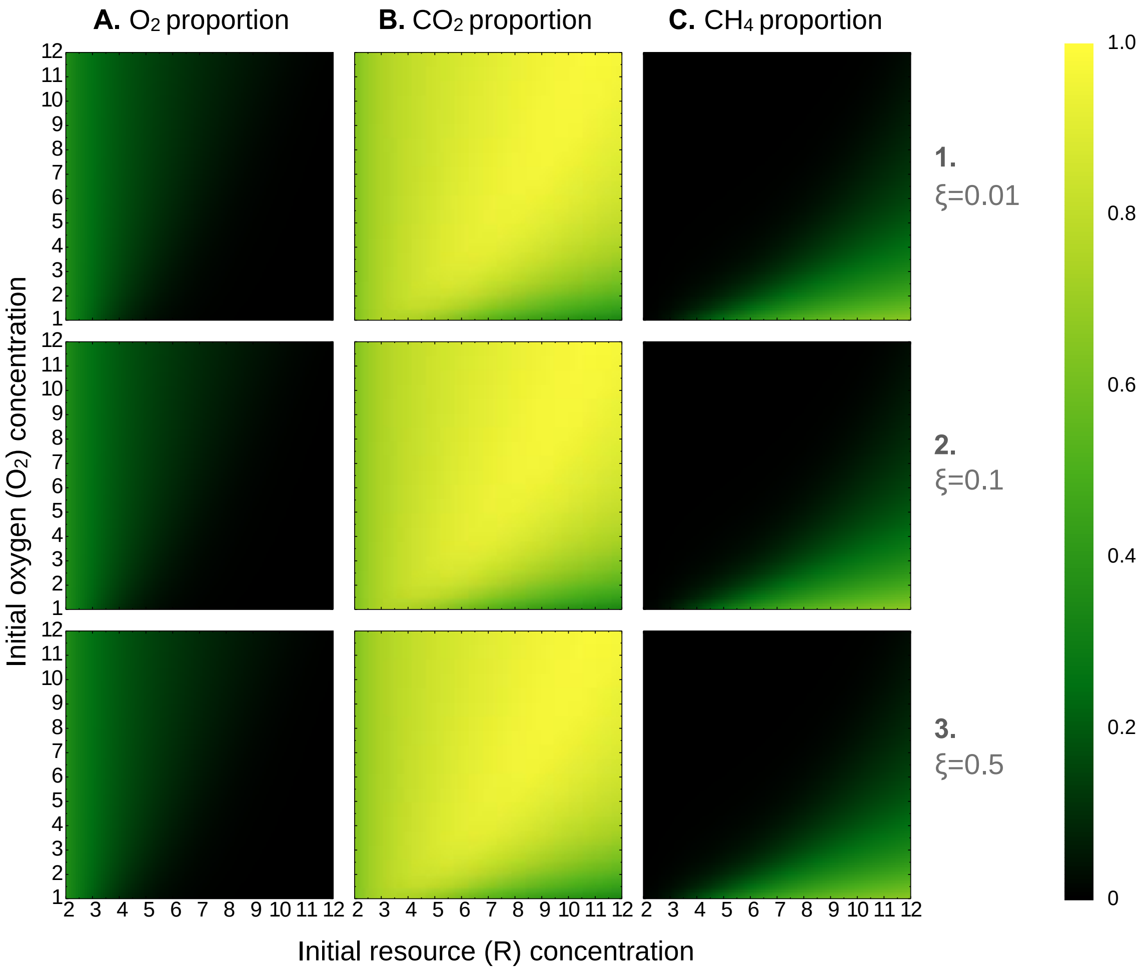


***Figure S4O:*** *Variations in $O_2$, $CO_2$, and $CH_4$ proportions (in columns A-C) run to steady state at set $O_2$ and R input values, with resource incorporation efficiency for fermenters (ξ) set at 1, 10, and 50% in rows 1-3, respectively.*

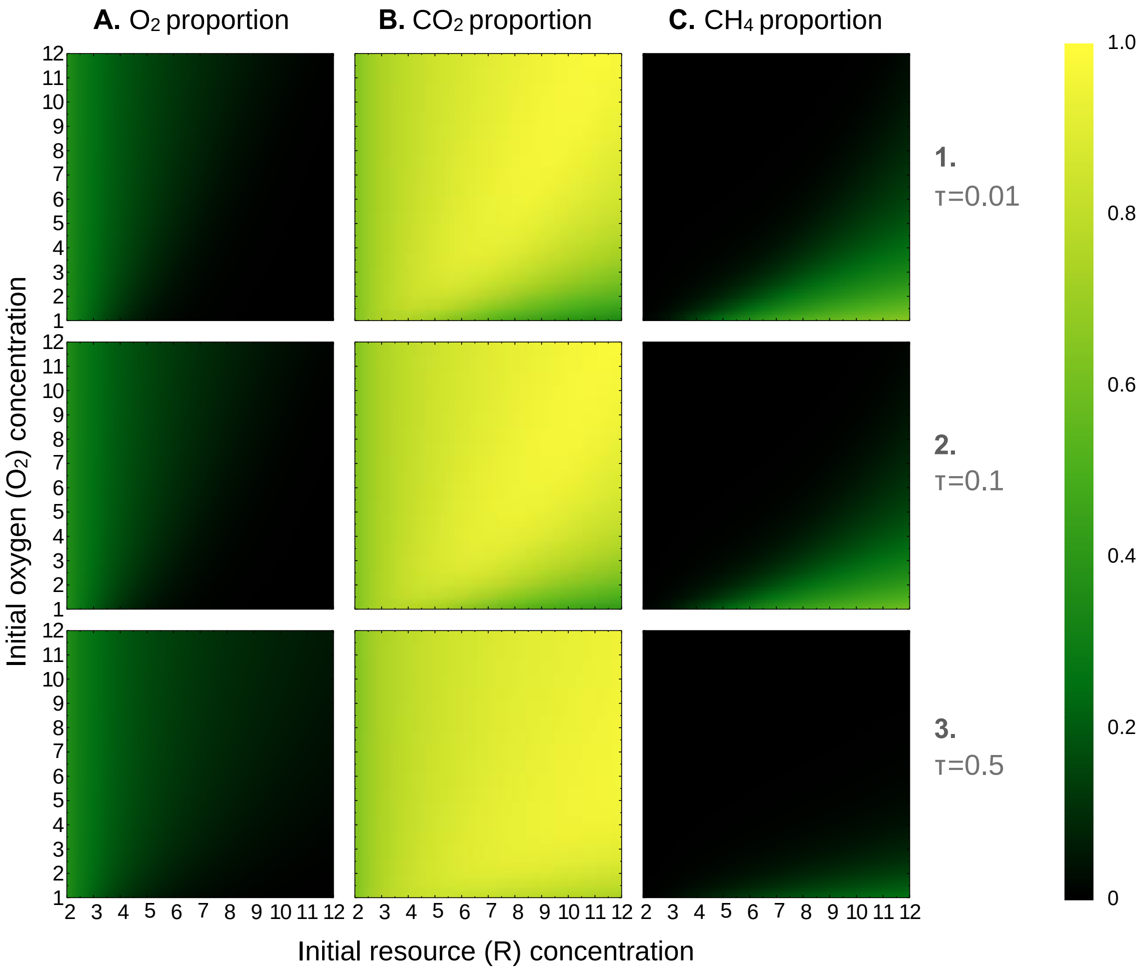


***Figure S4P:*** *Variations in $O_2$, $CO_2$, and $CH_4$ proportions (in columns A-C) run to steady state at set $O_2$ and R input values, with resource utilization efficiency for fermenters (τ) set at 1, 10, and 50% in rows 1-3, respectively.*

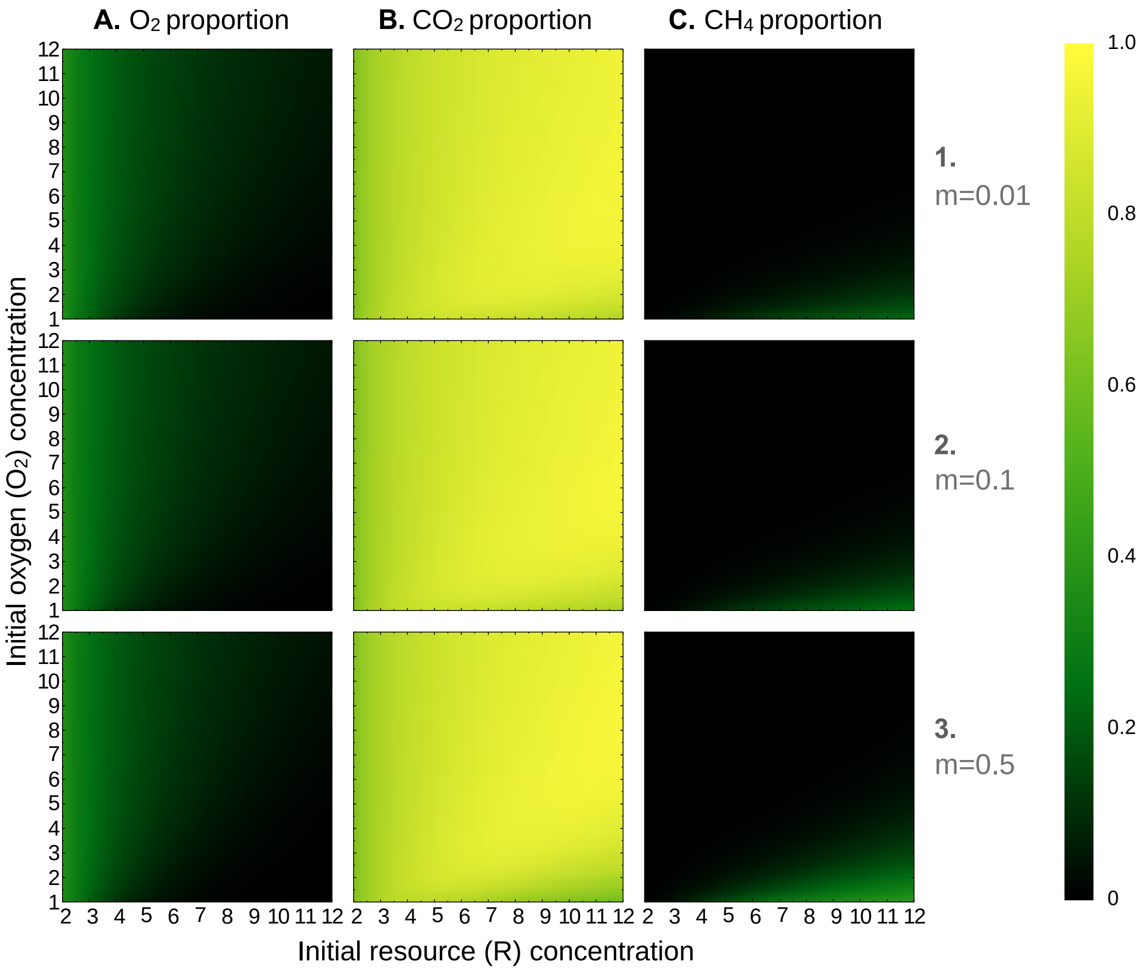


***Figure S4Q:*** *Variations in $O_2$, $CO_2$, and $CH_4$ proportions (in columns A-C) run to steady state at set $O_2$ and R input values, with energy requirement for anaerobic heterotrophs (m) set at 1, 10, and 50% in rows 1-3, respectively.*

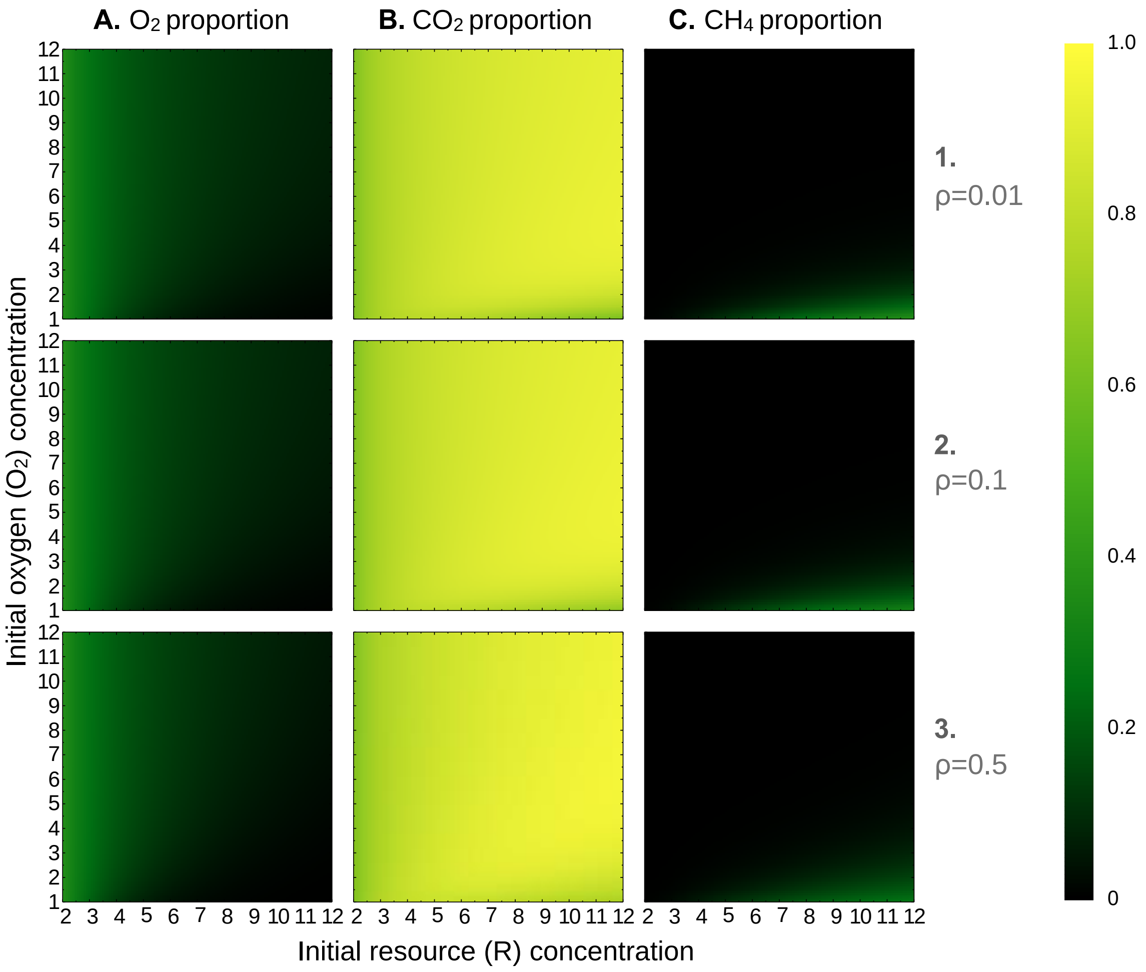


***Figure S4R:*** *Variations in $O_2$, $CO_2$, and $CH_4$ proportions (in columns A-C) run to steady state at set $O_2$ and R input values, with resource incorporation efficiency for anaerobic heterotrophs (ρ) set at 1, 10, and 50% in rows 1-3, respectively.*

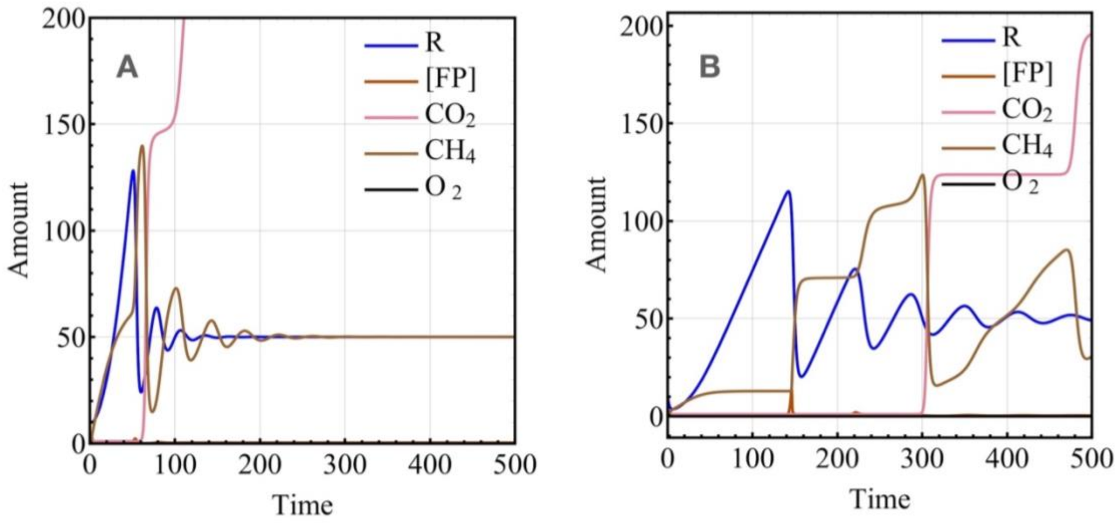


***Figure S5:*** *Long term dynamic variations in* $O_2$*, FP, and* $CH_4$ *run at two different initial* $O_2$ *concentrations, but constant input of R (resource) - (A) 5 units, (B) 1 unit. The initial* $O_2$ *concentrations affect the transient dynamics, but nevertheless the system converges towards the same stable state. These dynamics are reminiscent of the complex cyclic dynamics that we observe in the constant input of both resource and oxygen.*

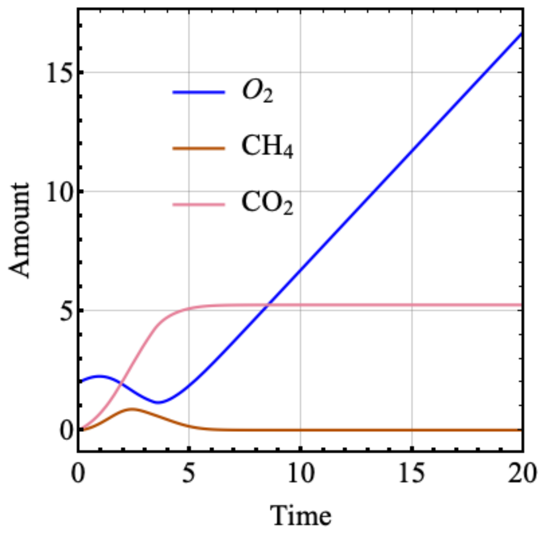


***Figure S6:*** *Dynamic variations in $O_2$, $CO_2$, and $CH_4$ under constant addition of $O_2$ and fixed initial amount of R (resource). In contrast to figure S5, due to lack of resources, different functional groups crash and the concentrations of gases/substrates (except oxygen) stabilize. Because oxygen is being pumped in continually, it keeps increasing.*

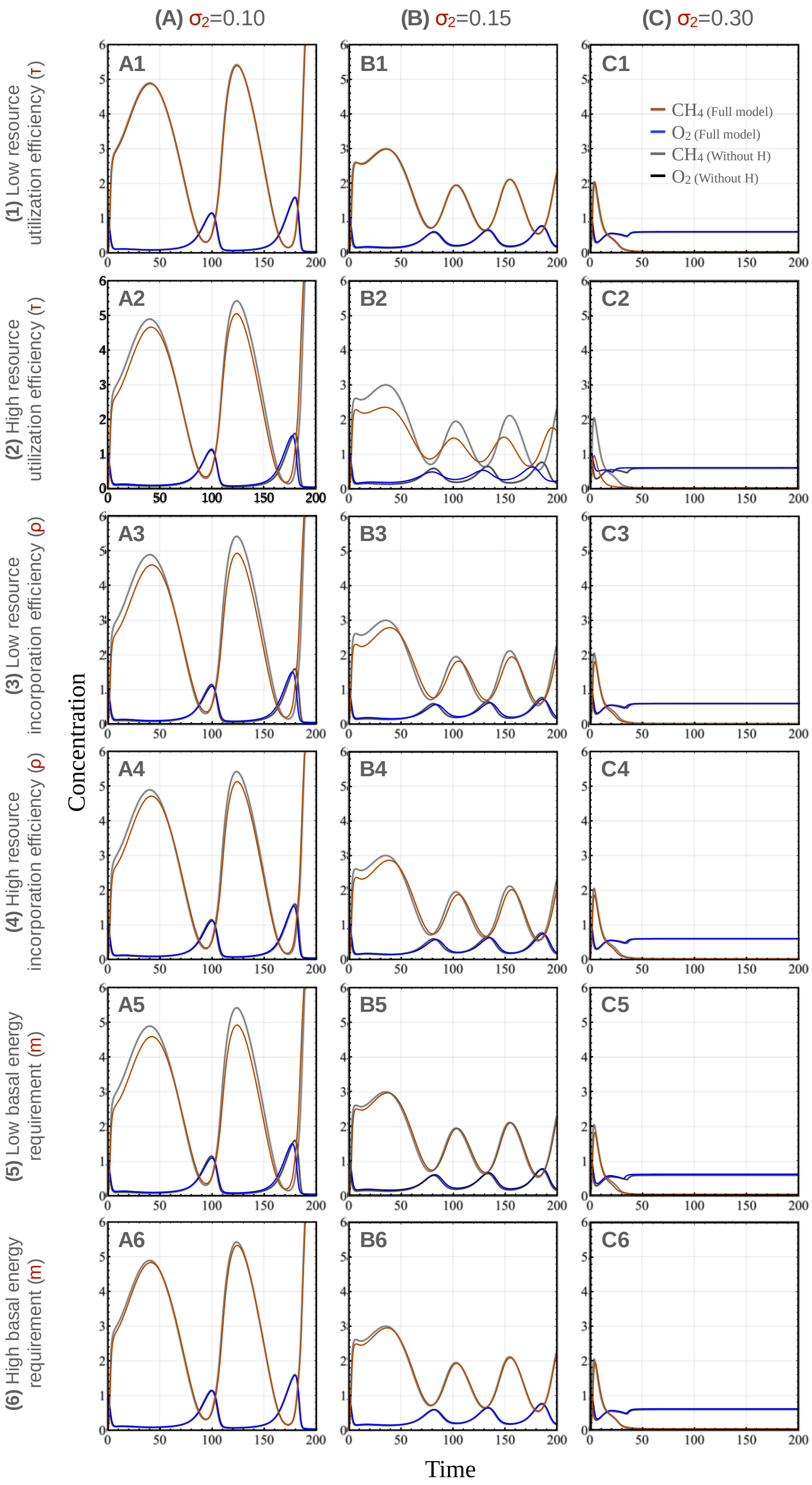
(A) σ2=0.10
(B) σ2=0.15
(C) σ2=0.30
(1) Low resource utilization efficiency (τ)
(2) High resource utilization efficiency (τ)
(3) Low resource incorporation efficiency (ρ)
(4) High resource incorporation efficiency (ρ)
(5) Low basal energy requirement (m)
(6) High basal energy requirement (m)
A1
B1
C1
A2
B2
C2
A3
B3
C3
A4
B4
C4
A5
B5
C5
A6
B6
C6
CH4 (Full model)
O2 (Full model)
CH4 (Without H)
O2 (Without H)
Concentration
Time

***Figure S7:*** *A comparison of concentrations of* $O_2$*, and* $CH_4$ *in time under three different regimes of addition of* $O_2$ *(*$\sigma_2$*) (and fixed rate of R (resource) addition (*$\sigma_1$*)) in the reduced model (full model without H) compared with the full model. The dynamics of methane and oxygen are similar in both the models and are mostly controlled by the relative addition rates of oxygen in both the models. This signifies the qualitative robustness of the model to addition of removal of new microbial functional groups (with respect to general behavior).*
